# The developmental gene *disco* regulates diel-niche evolution in adult moths

**DOI:** 10.1101/2023.05.28.542320

**Authors:** Yash Sondhi, Rebeccah L. Messcher, Anthony J. Bellantuano, Caroline G. Storer, Scott D. Cinel, R. Keating Godfrey, Deborah Glass, Ryan A. St Laurent, Chris A. Hamilton, Chandra Earl, Colin J. Brislawn, Ian J. Kitching, Seth M. Bybee, Jamie C. Theobald, Akito Y. Kawahara

## Abstract

Animals shift activity periods to reduce predation, minimize competition, or exploit new resources, and this can drive sensory system evolution. But adaptive mechanisms underlying niche-shifts are poorly understood, and model organisms are often too distantly related to reveal the genetic drivers. To address this, we examined expression patterns between two closely related silk moths that have undergone temporal niche divergence. We found 200-700 differentially expressed genes, including day upregulation in eye development and visual processing genes, and night upregulation of antennal and olfactory brain development genes. Further, clusters of circadian, sensory, and brain development genes co-expressed with diel-activity. In both species, eight genes showed expression significantly correlated to diel activity, and are involved in vision, olfaction, brain development, neural plasticity, energy utilization, and cellular maintenance. We repeatedly recovered *disco*, a zinc-finger transcription factor involved in antennal development, circadian activity, and optic lobe brain development in flies. While *disco* mutants have circadian arrhythmia, most studies attribute this to improper clock neuron development, not adult circadian maintenance. Comparing predicted 3D protein structure across moth and fly genetic models revealed *disco* likely retained developmental function with a conserved zinc finger domain, but gained functional zinc finger domains absent in *D. melanogaster.* These regions have several mutations between nocturnal and diurnal species that co-occur with higher levels of predicted phosphorylation sites. With robust circadian expression, functional nocturnal and diurnal mutations, and structural and sequence conservation, we hypothesize that *disco* may be a master regulator contributing to diel-activity diversification in adult moths.

**Significance:** Insect diel-activity patterns are diverse, yet the underlying evolutionary processes are poorly understood. Light environment powerfully entrains circadian rhythms and drives diel-niche and sensory evolution. To investigate its impact, we compared gene expression in closely related day- and night-active wild silk moths, with otherwise similar ecologies. Expression patterns that varied with diel activity included genes linked to eye development, neural plasticity and cellular maintenance. Notably, *disco*, which encodes a zinc-finger transcription factor involved in pupal *Drosophila* optic lobe and antennal development, shows robust adult circadian mRNA cycling in moth heads, is highly conserved in moths, and has additional zinc-finger domains with specific nocturnal and diurnal mutations. We hypothesize that *disco* may contribute to diversification of adult diel-activity patterns in moths.

## Introduction

Circadian rhythms regulate many biological processes, including day-night activity patterns. Research to date has explored genes, circuits, and environmental cues mostly in the context of control within a single organism (1, 2). Diel-niche stems from the concept of temporal partitioning of activity periods, driven by the variation in resources, temperature, light level, or predation, across the day (3–6). Periods are often binned into three distinct groups: diurnal (day active), nocturnal (night active), or crepuscular (active at dusk, dawn, or both), and although this is useful for comparative analysis, it is often an overgeneralization (7–11), and the evolution of diverse diel-niches across organisms (7, 11–15) is poorly understood (16–19).

Several studies have documented sensory adaptation accompanying evolutionary diel-niche transitions in mammals, birds, and insects (8, 18, 20, 21). Specific examples include colour vision gene expansions in diurnal moths (22, 23), improved olfactory senses in dark-bred flies (24), and the evolution of hearing organs in nocturnal butterflies (25). Comparisons of circadian genes across model organisms catalogue variation at long evolutionary timescales and highlight conserved elements (26, 27), but diel switches can occur over short timescales (28), and distantly related species provide less insight into mechanisms that enable faster switches. Moths and butterflies (Lepidoptera) are an ideal group to address this, as their evolutionary history is well known and there are many diel-niche switches, often between closely related species (7, 29). They are also one of the few insect groups, outside of *Drosophila*, in which both sensory and circadian genes have been characterized (23, 30–35).

Wild silk moths (Saturniidae) are important models for understanding chronobiology, with seminal experiments confirming the brain functions as the primary circadian control (36, 37). While most are nocturnal, many fly during the day, and temporal niche switches may function as a mechanism for reproductive isolation in sympatric species (38, 39). Saturniid phylogeny is relatively well known, and there are several annotated genomes, chromosomal maps. Furthermore, they belong to the superfamily Bombycoidea, which includes the domesticated silk moth, *Bombyx mori*, a key biological model organism serving as a reference taxon for comparative genomic studies (32, 40–43). Saturniids such as *Antheraea* are of major importance to the silk industry, and they have a well-resolved taxonomy and well-documented life histories (38, 44–46). Despite this, the genetic elements that control day-night activity and diel-niche evolution, in this group and insects in general, remain largely unknown (28, 37).

We used RNA-Seq to characterize expression patterns during peaks and troughs (midday and midnight) across two closely related wild silk moths: the diurnal *Anisota pellucida*, and nocturnal *Dryocampa rubicunda*. They are sister genera, feeding on large deciduous trees in the temperate forests in Eastern North America, with a recent divergence in temporal niche ∼3.8 Mya (47). We sequenced head tissue at different time periods to identify expression in sensory, circadian, and neural genes that correlate with diel activity. Expression in eight genes clearly and significantly correlates with diel activity, with functions in vision, olfaction, brain development, neural plasticity, energy utilization, and cellular maintenance. Of these, a single gene emerges consistently across analyses: *disco*, which encodes a zinc-finger transcription factor. In moths, it has likely retained functions also found in flies (eye and brain development during the pupal stage) through a conserved zinc-finger domain. But it has also gained an extra zinc-finger domain, surrounded by phosphorylated sites and with mutations in both nocturnal and diurnal species, which may contribute to adult moth circadian regulation.

## Results

We compared gene expression across two wild silk moth species, *Anisota pellucida* and *Dryocampa rubicunda*, whose males are diurnal and nocturnal, respectively (Fig. 1, Table S1). We generated head (eyes, brain) transcriptomes from moths collected and flash frozen at midday and midnight, referred to as ‘day’ and ‘night’ hereafter. Using multiple programs to assemble high-quality *de novo* assemblies (Table S2), we characterized the level of gene (mRNA) expression with quasi-read mapping, which can be used to correlate protein expression (48).

**Figure 1:**
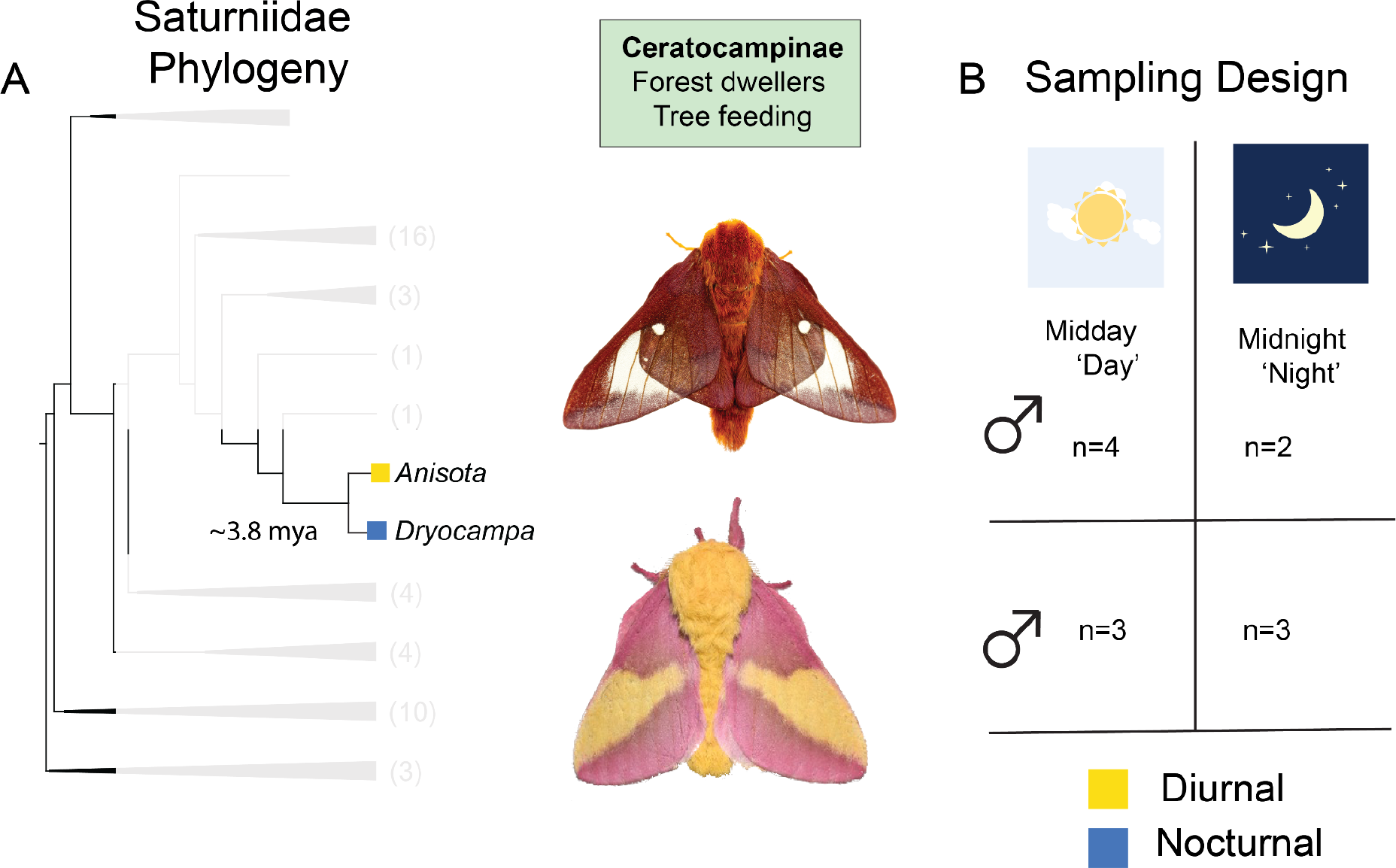
Nocturnal and diurnal moths on a phylogeny with RNA-seq sampling design. **(A)**: Collapsed phylogeny of Saturniidae, adapted to show where the two study species, diurnal *Anisota pellucida* (Pink-striped oakworm moth) and nocturnal *Dryocampa rubicunda* (Rosy Maple moth) sit. Faded numbers on the tip represent the number of genera in the tree before collapsing. Phylogeny adapted from Rougerie et al. 2022 (**B)**: Sampling design showing the number of replicates for each species and collection period (day/night) sampled. Collection of heads was done 2-days post eclosion at midday (sun) and midnight (moon) and tissue was flash frozen for RNA preservation. Care was taken that the eclosed moths were exposed to a natural light cycle and red lights were used when collecting the moths at night. Photo credits *Anisota pellucida* © Mike Chapman; *Dryocampa rubicunda* (CC) Andy Reacgo and Chrissy Mclearan;

### Day-night gene expression patterns switch between nocturnal and diurnal species

We found 350 & 393 significantly differentially expressed genes (DEGs) when comparing day and night treatments for each species (Table S3, Fig. 2A, Supplementary Data 1). *Anisota* had more day- upregulated genes (56%) and *Dryocampa* was slightly more night-upregulated (53%). In order to compare DEG sets between species, we mapped our DEGs to *Bombyx mori* orthologs using the Orthofinder software (49) (Supplementary Data 1). Approximately 60% of DEGs from each species had identifiable orthologs in *B. mori* (Table S3), and only a small number of DEGs (6-8 genes) overlapped between both species (Fig. 2B). We also replicated this analysis using other software to assure that our results were robust, despite different normalization methods (50). With DESeq2, we found 498 & 697 DEGs (Fig. 2C, Table S3, Supplementary Data 2), with similar *Bombyx* annotation rates (61%), although the proportion of day upregulated genes increased considerably in the *Anisota* (79%) compared to being more even split in *Dryocampa* (50%) (Table S3). The total number of overlapping genes increased (19-26, Fig. 2D) when using DESeq2. A comparison of the two methods revealed that 174 and 216 genes were shared between *Anisota* and *Dryocampa*, respectively.

**Figure 2:**
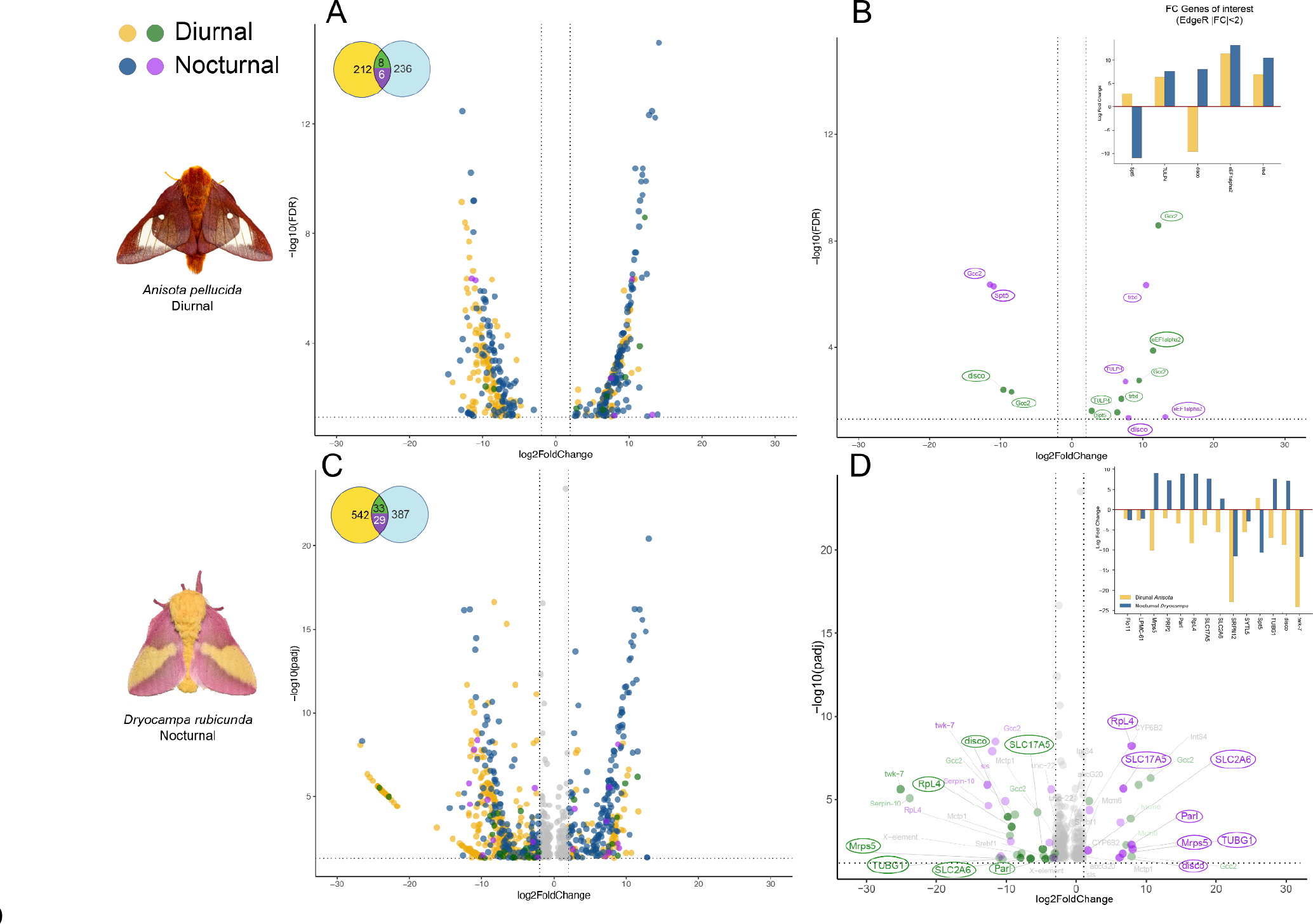
Nocturnal and diurnal species show divergent patterns of gene expression across two different analyses software (EdgeR and DESeq2) Volcano plots and Venn diagrams showing EdgeR (A-B) and DESeq2 (C-D) results across **day vs. night** sampling times for both nocturnal and diurnal species. Venn diagrams represent the number of differentially expressed genes (DEGs) across both species with the number of common DEGs across each pair. **(A/C)**: Volcano diagrams illustrating fold change and adjusted p-values for the significant differentially expressed genes between midday and midnight samples. Circles in the top left represent the number of genes expressed in both species and the colors correspond to FC values for those genes in the volcano plots. Yellow and blue represent genes expressed only in the nocturnal or diurnal species, green/purple indicates DEGs present in both species. **(B/D)**: Only genes expressed in both species are shown and genes that display opposite trends in expression are highlighted. See Supplementary Material for a detailed list of DEGs. Positive fold change indicates night overexpression and negative fold change indicates day overexpression. Genes had FDR or adjusted p-value < 0.05 and FC > |2|. Identification of common genes was done using orthogroup clustering with Orthofinder with *Bombyx mori*. Gene names and annotations were transferred from *B. mori*.

### Divergently expressed genes are linked to brain optic lobe, antennal and neural development

To identify important regulators involved in diel-niche evolution, we applied two filtering criteria to our gene expression data. First, we selected genes that exhibited highly significant differential expression in both species. Second, we focused on genes that displayed upregulation patterns consistent with the natural diel-activity of each species. Our rationale was that this subset of genes was more likely to contain key regulators. To compare DEG overlap between the two species, we grouped the transcripts to their matching orthologs from *Bombyx mori*; if two transcripts from different species mapped to the same ortholog, we treated them as being the same. This allowed us to examine overlapping genes between the species to see if any genes switched fold-change sign from positive to negative or vice versa. (Fig. 2, Supplementary Data 3). We found 51 overlapping DEG transcripts that mapped to 28 unique *Bombyx* genes. Nine genes showed flipped patterns of expression between the two species, and eight of them coincided with known diel activity patterns (Table S7). Examining gene ontology (GO) annotations and comparing orthologs from flybase (https://flybase.org/), we found genes linked to optic lobe and antennal development (*disco*), locomotion and energy use (*SLC2A6, SLC17A5*), brain and neural development (*TUBG1*) and other essential biological processes like transcription, ribosomal translation, protein processing, mitochondrial maintenance (*RpS4, PARL, Mrps5*) and wound response (*PRP2*) (Table S7, Fig. 2B, D). Of these, only *disco* was recovered with both methods.

### Gene network analysis identifies diel activity and species-specific co-expressed clusters

Identifying highly expressed genes helps understand which genes are activated during particular biological processes. However, determining only those that are highly expressed can often overlook genes with important biological functions (51). We examined co-expressed genes that may be correlated with diel-niche or RNA collection time. We used WGCNA, a weighted correlation network analysis tool to cluster genes together based on their normalized counts (52). After examining co-expression patterns for each species separately, we found two modules in each species that clustered with day-night treatment (cluster-*grey60* and cluster-*tan*) (Fig. S6, Supplementary data 4). Since we were interested in species-specific differences, we reran analyses and combined reads from *Anisota* and *Dryocampa,* using only normalized counts for genes that had valid *Bombyx* annotations for both species. This approach narrowed our focus to 2000 genes. Among these, we discovered two clusters (cluster-blue and cluster-turquoise) consisting of 50 genes each that exhibited different expression patterns across species (Fig. 4).

### GO enrichment of photoperiodism, circadian control, muscle and neural growth genes

We used a gene enrichment analysis to determine if gene ontology (GO) terms were significantly overrepresented in the DEG and WGCNA sets compared to the appropriate background of GO terms. Using TopGo, which allows custom gene sets, we found an overrepresentation of genes involved in several biological functions (Supplementary Data 5). Those that seemed significant for diel-niche and vision were ‘response to stimulus’ and ‘smooth muscle control’, as well as folic acid serine, glycine and retinoic acid metabolism. We also used ShinyGo to examine WGCNA clusters (Supplementary Data 5). We matched orthologous genes in the less duplicated, filtered transcriptomes to obtain identifications of the most related *Bombyx* genes (Table S4). We examined the enrichment of both tan and grey60 modules, listing the non-redundant terms using ReviGo (Fig. S7, Supplementary Data 5). Gene clusters that co-expressed in the same direction together in the day and night treatments of both species included photoperiodism, circadian control, negative phototaxis and nervous system development. We next checked for the enrichment of the modules that showed species-specific patterns (blue and turquoise, Fig. S8, Supplementary Data 5). These included genes involved in muscle proliferation and nerve growth, neural signaling, glycolysis, oxidative stress response, and basic cellular functioning such as protein processing and transcriptional regulation.

### Day upregulation of vision genes in the diurnal moth

Since both EdgeR and DESeq2 analyses use different normalization methods and statistical model assumptions (50, 53), we repeated enrichment analyses by combining datasets and examining genes that appeared in both analyses. For the *Anisota,* we tested over-enrichment of a smaller subset (FC ≤ -5) of diurnally highly upregulated genes (Fig. 3, Table S5). We found gene enrichment for visual perception, excretion regulation, negative gravitaxis, synaptic plasticity, along with genes associated with other biological processes, such as RNA interference, endopeptidase activity and endocytosis. A reduction in stringency (FC ≤- 2) did not alter results considerably (Fig. S4). Night-upregulated genes (FC ≥ 2) included ocellar pigment genes, eye-photoreceptor cell development, snRNA processing, post embryonic development and neurotransmitter secretion, among a host of other processes that may be required for growth, development and metabolism (wnt signaling, tricarboxylic acid cycle, and cellular response to insulin, glucose transport) (Fig. S4).

**Figure 3:**
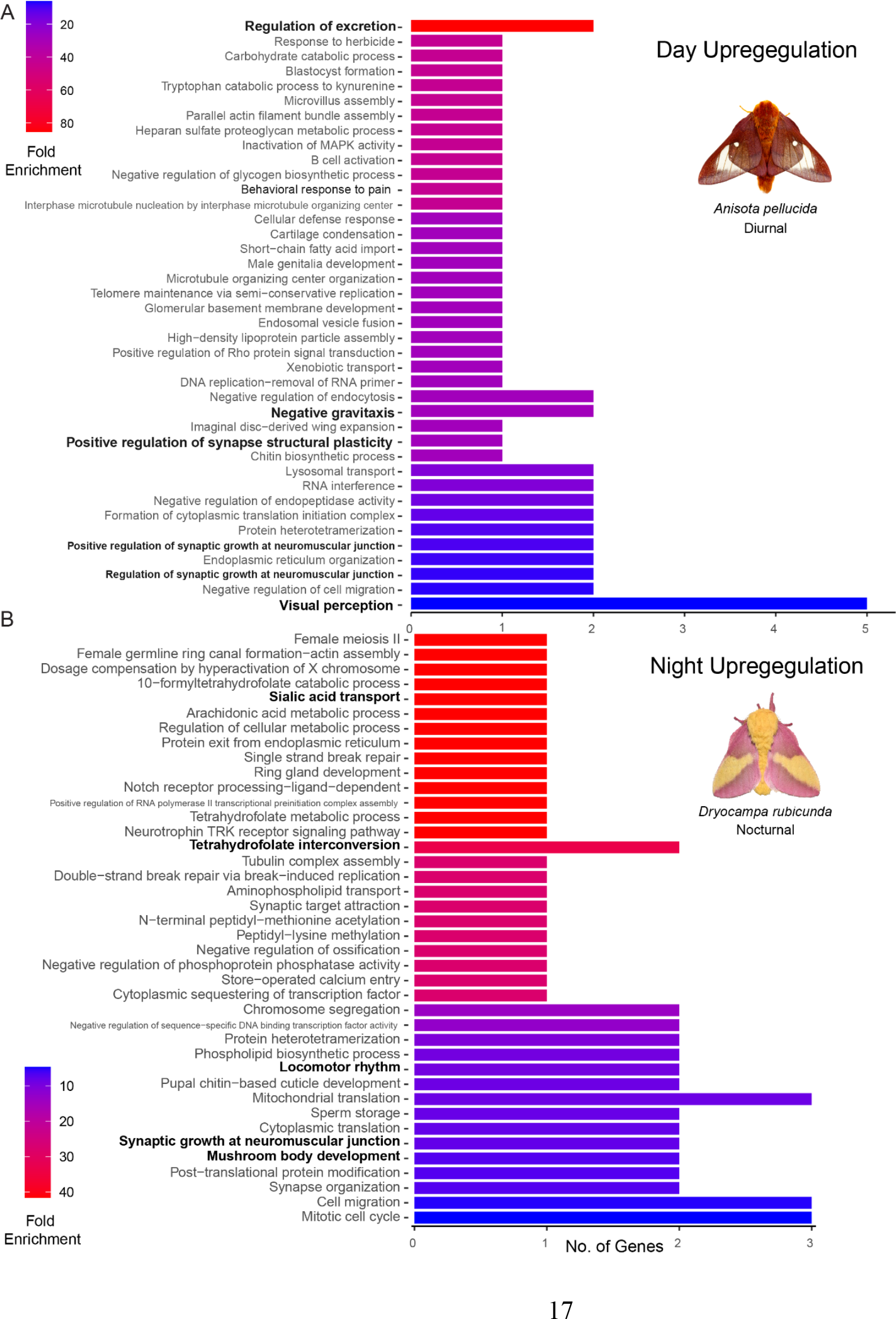
Visual genes are upregulated in the diurnal species during the day, and energy utilization, brain olfactory region and locomotion in the nocturnal species during the night. Go enrichment of highly upregulated genes recovered from both DEG analyses coinciding with the species. highlighting the two modules that showed species specific clustering patterns. A: Enrichment of **day** upregulated genes in diurnal *Dryocampa*. B Enrichment of **night** upregulated genes in nocturnal *Anisota.* Go enrichment was done using the custom ShinyGo database v0.75c using *Bombyx* gene IDs. FDR cutoff <=0.17, and only Biological Process GO terms were selected with Min. pathway size =1. Input genes had FC>5, padj. <0.05 and recovered both in EdgeR and DeSeq2 were used with the background being all orthologous *Bombyx mori* genes for either species.

### Night upregulation of antennal and olfactory brain regions mushroom development genes

We repeated the same analyses for *Dryocampa* and tested highly nocturnally upregulated genes (FC ≥ 5). Our results show upregulation in genes known to be associated with mushroom body development, locomotor rhythm, synaptic growth, energy utilization (Sialin transport) and mitochondrial translation (Fig. 3B, Table S6). A reduction in stringency (|FC| ≥ 2) showed entrainment of the clock cycle, and antennal development genes. Genes associated with innate immune response, DNA repair, cell division, histone acetylation, circadian rhythm, retinoid cycle were upregulated during the day, possibly indicating a period of cellular repair during a time when these moths are inactive (Fig. S5).

### Key sensory, circadian, eye development and behavioral genes can drive diel-niche switches

We combined results from the DEG (EdgeR and DESeq2) and gene network analyses (WGCNA) to create a cumulative list of 1700 transcripts (Supplementary Data 6). Focusing on genes that were recovered across *Anisota* and *Dryocampa* reduced the set to 274 transcripts (Supplementary Data 7). Because many transcripts had poorly annotated *Bombyx* hits, we improved annotations using the program eggNog mapper (54). We tested if these genes had GO terms associated with sensory, circadian, brain and neural development, or behavioral regulatory genes (Fig. 4, Table S7, Supplementary Data 8). We found that several genes in each category had associated GO terms, with a predominance of vision and brain development genes (Supplementary Data 9).

**Figure 4:**
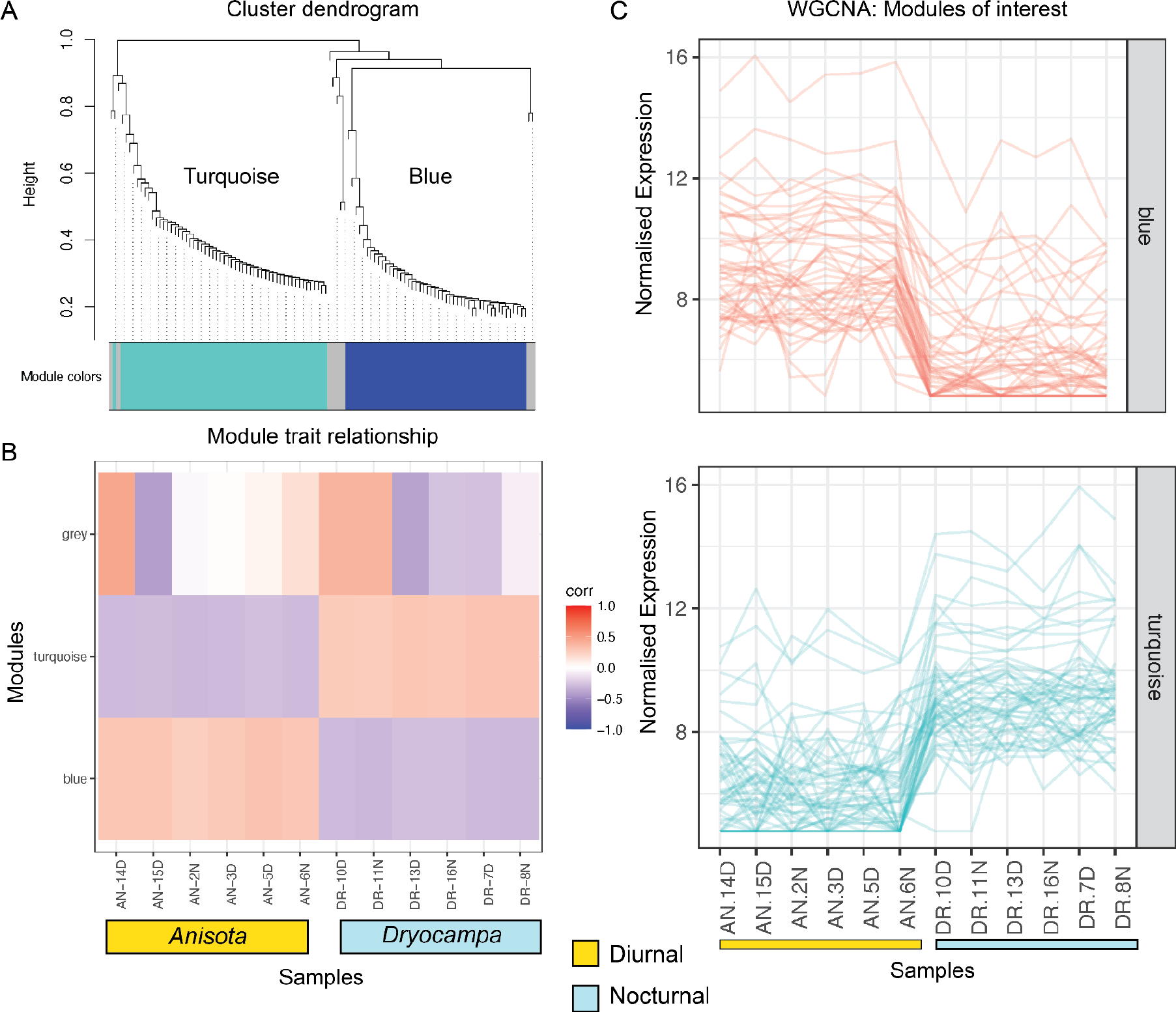
Modules of clustered co-expressed genes grouped using normalized expression highlighting the two modules that showed species specific clustering patterns. A: Cluster dendrogram shows WGCNA clusters. B: Shows how patterns of gene expression correlate across samples and modules. C: Shows the normalized expression for all genes across samples. The species show two sets of 50 genes (Blue) and (Turquoise) that have clear species specific expression patterns. Normalization was done with DESeq2 and reads were mapped to the more stringently filtered transcriptome. A soft power analysis was done and the picked power=9 for the WGCNA analysis. AN: *Anisota*, DR: *Dryocampa,* Numbered D and N represent the different samples collected at day and night time points.

**Figure 5:**
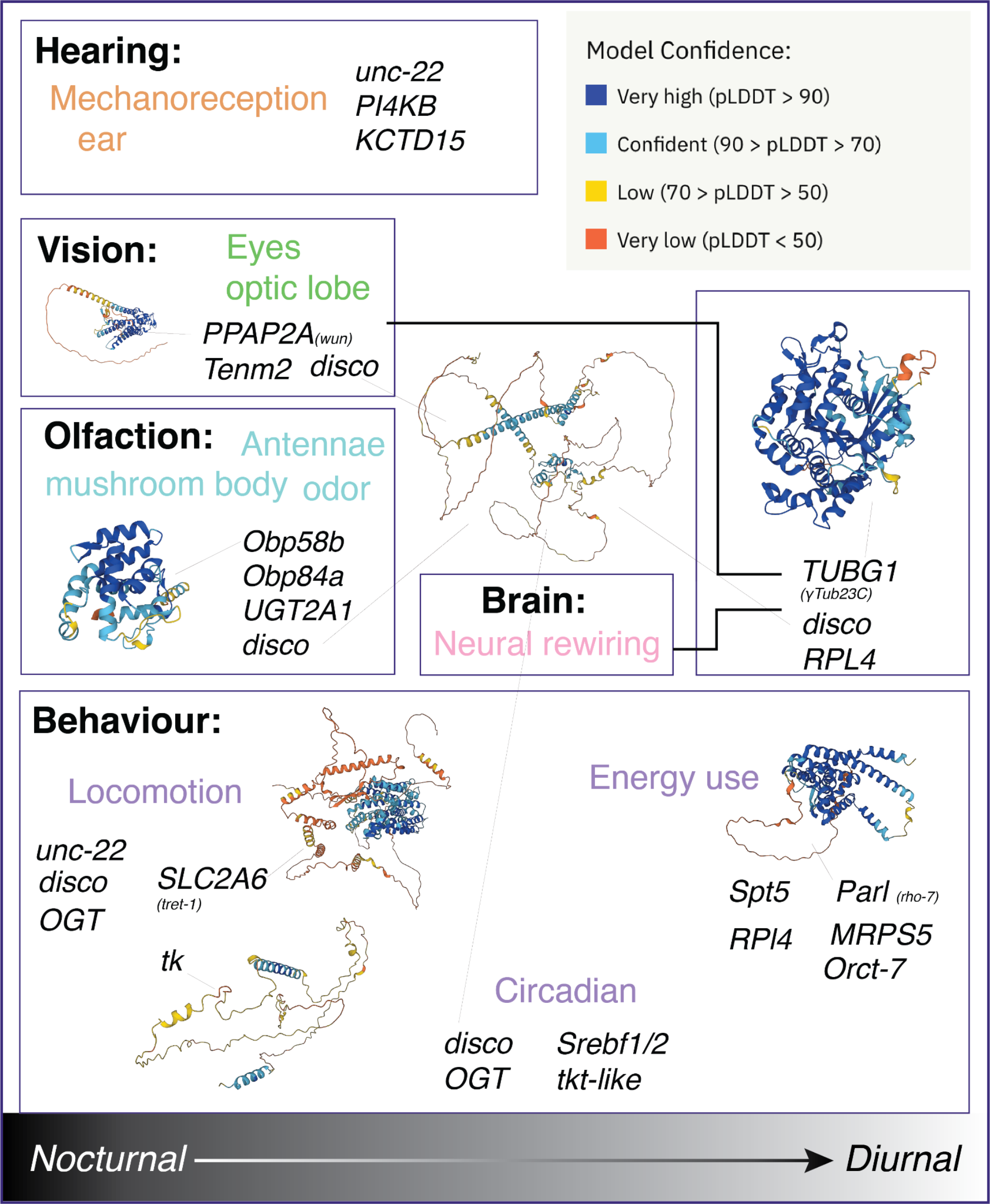
Model for diel-niche shift from and genes of interest obtained from GO searches and annotating divergently expressed genes. For representative purposes, homologs from *Drosophila melanogaster* have been used, with colors in the 3d structures representing Alpha fold per-residue confidence metric (pLDDT), the range for each color is shown on the top left. Model moth *(Bombyx mori)* proteins were also modeled for a subset of proteins, see Supplementary data.

### Predicted functional regions and homology patterns identified for genes of interest

We examined protein and gene evolution for a set of genes which we found were of interest based on results from DGE, WGCNA, GO annotations (Supplementary Data 10). In order to infer functional homology from structural conservation, we downloaded high quality genomes of Bombycoidea moths and relatives from Darwin Tree of Life [https://www.darwintreeoflife.org/] (Supplementary Data 11). We assigned a reference protein sequence for each gene of interest from the *Bombyx mori* predicted proteome. We used Orthofinder to identify orthologs and filtered orthogroups containing the reference sequence (49, 55). Since we had more than one gene per species, we reduced the number down to a single gene by choosing the highest identity sequence relative to the *B. mori* reference sequence. We also modelled the 3D structure of *B. mori* proteins and mapped the evolutionary conservation onto the 3D predicted structure for proteins above a certain conservation threshold (Supplementary Data 12). These analyses predict structurally and functional conserved regions of proteins (Fig S9, Supplementary Data 13). We repeated this analysis with 38 insect genomes (Supplemental Data 11) and mapped evolutionary conservation onto 3D protein structure (Supplementary Data 12-13). We include results of evolutionary conservation analyses for two regulatory candidates (*disco* and *tk*) that showed varying levels of sequence and protein evolution between insects and moths (Fig. S10).

### Modeling predicts additional functional zinc-finger domains for disco in Lepidoptera

*Disco* was recovered across multiple analyses. In *Drosophila melanogaster* it is known for its role in eye development, important for circadian maintenance, and for leg and antennal appendage formation (56–60). To determine if *disco* was conserved between moths and *Drosophila*, we compared the primary sequence and 3D protein structure of *Drosophila* and *Bombyx mori*. In the moth, the sequence length of *disco* was nearly double that of *Drosophila* (Fig 6A). However, a region spanning over 100 amino acids was highly conserved, contained the zinc-finger domain important for its function, and showed strong 3D structural conservation (*Whole protein alignment*: RMSD: align, super) = 37.833, 2.566, MatchAlign score (align, super) = 540 (2234 atoms), 333.1 (554 atoms) vs. *alignment of conserved region*: RMSD=2.699,2.566, MatchAlignScore =443(671 atoms), 351 (615) atoms). This indicates the DNA binding function of *disco* has likely been conserved. However, an additional ∼500 amino acid region absent in *Drosophila* is highly conserved across moths, and includes several regions predicted to be functional (Fig. 6A, Fig. S10). We hypothesize that *disco* has a novel role in moths for diel-niche regulation. To further test this, we compared *disco* sequences across *Anisota* and *Dryocampa* and found 23 mutations between them, three of which mapped to the predicted functional region (Fig. 6B, Supplementary Data 14). InterProScan predicted four zinc-finger domains, three in this region, although the CATH-Gene3D databases prediction combined the two separate domains into a single predicted domain (Fig. S11). We also found 53 sites with predicted phosphorylation potential, especially around the second zinc-finger domain (18). Reducing the stringency increased the total to 142, which were still enriched around the second domain.

**Figure 6:**
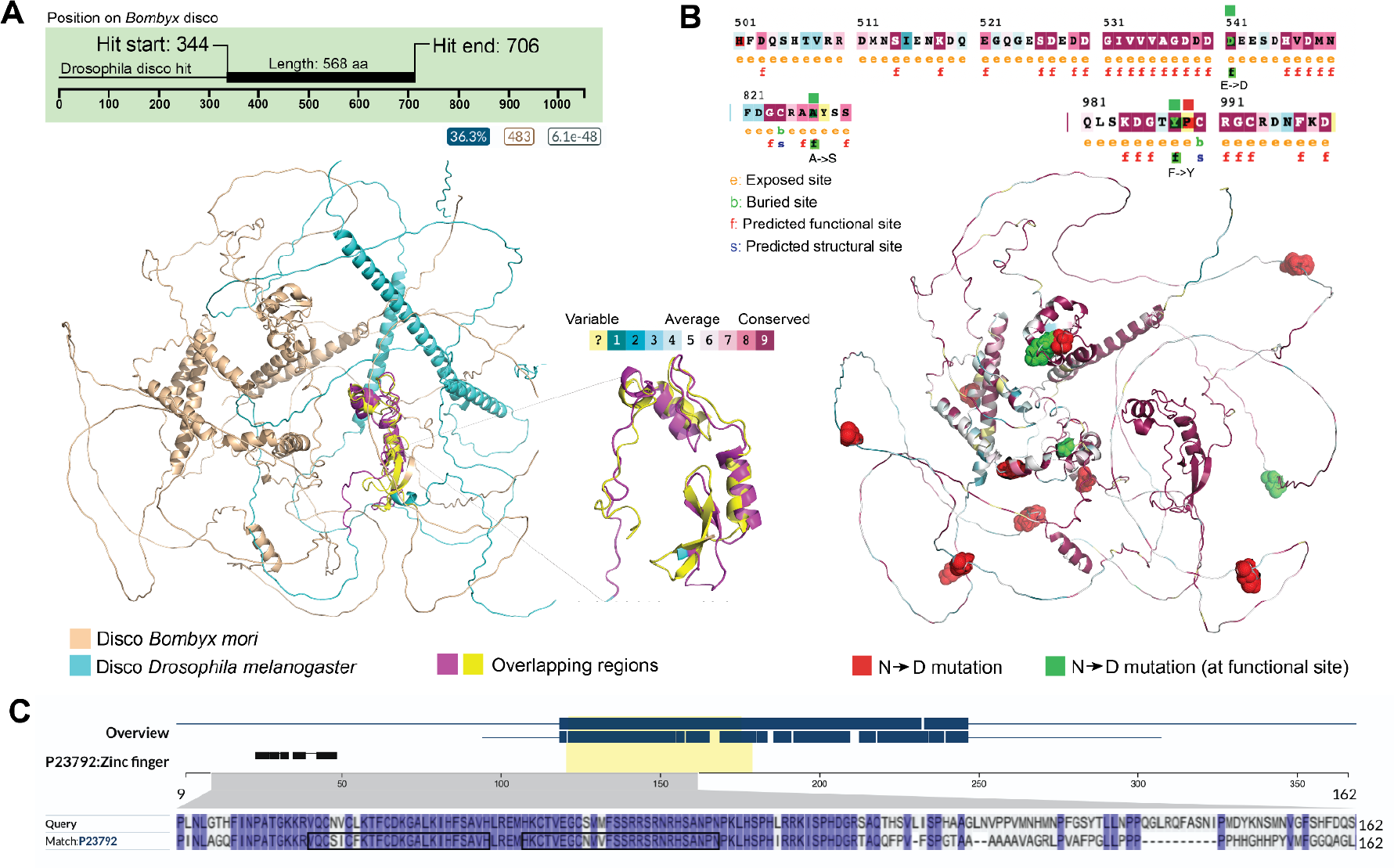
*Disco* has 150 amino acid long highly conserved region across *Bombyx* (1054 aa) & *Drosophila* (568 aa) involved in its role as a transcription factor, but it also has other roles in Lepidoptera due to the presence of other highly conserved and predicted functional regions, that are not present in flies, and only in Lepidoptera. 3/14 of the mutations between nocturnal to diurnal species map to these regions (>500 aa). **(A)** Top: *Disco* best Uniprot hit in *Drosophila* using default settings (blastp, e-threshold 10, Auto- Blosum62). Bottom: *disco* aligned and superimposed predicted structures for the silk moth and vinegar fly. Peach structure: *Bombyx mori* alpha-fold predicted structure for *disco,* Cyan: *Drosophila melanogaster* alpha-fold predicted structure for *disco* **(B)** Top: Partial views of the Consurf predicted residues for *disco.* The views encompassing the region where 3 mutations of the 14 mutated sites between Nocturnal *Dryocampa* and Diurnal *Anisota* overlap with predicted functional residues. Bottom: Residues that were mutated between the nocturnal and diurnal species are highlighted on an overlay of the Consurf scores mapped onto the predicted alpha fold structure of disco from *Bombyx mori.* Green residues indicate mutated and predicted functional sites, red indicate mutated sites that did not have a predicted residue. **(C)** Overlap of the highly conserved region of *disco* this region includes the zinc-finger domain that is characteristic of disco transcription factor.

## Discussion

We identified the genetic mechanisms of diel-niche switches by examining gene expression during the day and night in two closely related moth species. We used RNA-Seq to measure gene expression profiles of head tissue and isolated sensory, circadian and neural processes. We found between 300 and 700 genes that significantly altered day night circadian expression patterns. The diurnal species had enriched visual perception genes during the day and the nocturnal species had locomotor and olfactory genes upregulated at night.

Thirty overlapping DEGs were present in both species, with some DGEs showing divergent patterns of expression matching the species’ diel-niche. Examining expression data with a sensitive clustering analysis yielded over 170 genes that showed day-night or species specific co-expressed clusters. We also used GO terms to search for genes associated with sensory, neural, or circadian patterns. We found genes in each category and modeled the 3D structure and predicted functional regions for a subset. The divergently expressed gene *disconnected* (*disco*) was implicated in vision, hearing, locomotion and brain development. Given the multiple lines of evidence supporting the importance of this gene in regulating diurnal and nocturnal activity in moths, we explored the evolution of *disco* by modeling fly and moth *disco* structures. This analysis revealed novel zinc-finger conserved domains in moths, which were lacking in *Drosophila,* surrounded by phosphorylated sites. Several mutations between *Anisota* and *Dryocampa* also mapped to these regions, further strengthening evidence for its role in diel-niche shifts.

### Visual and olfaction

Visual systems often accompany diel and photic environment shifts (18). For example, nocturnal carpenter bees have much larger facets in their eyes than their diurnal counterparts (61). Nocturnal moths have higher photoreceptor light sensitivity and have neurons more suited to pooling than close diurnal relatives (62). Some of these trends seem to hold across the phylogeny, with diurnal species evolving more complex color visual systems, often reflected in visual opsin sequence evolution (22, 23). We did not find diel-expression patterns in color vision opsins, a result corroborated by a recent study (22). However, we discovered a cerebral opsin (ceropsin) upregulated during the day. Ceropsin has been implicated in photoperiodism and is expressed in the brain (63).

We also found several eye development genes (*TENM2, ANKRD17, EHD4, JAK2*), phototransduction genes (*PPAP2*, *RDH11*), and retina homeostasis, eye-antennal disc development, and photoreceptor cell maintenance genes (*disco, glass*) (64). Surprisingly a few visual genes, such as *garnet* and *rugose* also appeared to have different isoforms present, showing both day and night upregulation. *Garnet* is an eye color mutant gene in flies (65) and *rugose* is implicated in retinal pattern formation (66).

One of the classical differences between day flying butterflies and night flying moths is that butterflies have a clubbed antennae, while moths often have a sensilla-covered antenna; likely a function of increased investment in olfactory sensing, similarly, there is also evidence for increased investment in olfactory sensing in the brain, with relatively larger mushroom bodies in nocturnal species (67, 68). In our diel gene-expression dataset, we found several olfactory genes, including those involved in odorant binding (*Obp84a, Obp58b*), pheromone response (*tk*), mushroom body development (*DAAM2 and DST*) and antennal development (*disco*).

### Hearing and mechanoreception

Many moths have active hearing organs, and these have evolved repeatedly across Lepidoptera (69), and their use varies depending on the ecological context of the moth. Several drivers for moth hearing include sexual selection and as a defense against insectivorous bats (70). The primary nocturnal group of butterflies, Hedylidae, has reverted to a nocturnal niche and regained hearing organs (25). Thus, although rudimentary vibrational hearing exists in some diurnal moths (71) it is significantly more developed and used at broader frequency ranges at night, when light is less available. While Saturniidae lack hearing organs (72) they may instead still have vibration mechanosensors that can be useful in evading predation similar to those in locusts and crickets (73–76). Indeed, we did find diel co-expression of several mechanosensory and ear development genes, including *PI4KB, KCTD15, unc-22,* and *RhoGAP92B* (77)

### Brain and neural rewiring

Finding an upregulation of neural and brain development was expected since we sequenced head transcriptomes. However, we tested gene expression two days after pupal eclosion, where presumably gross structures are already developed, so it is possible these expressed genes we found, regulate adult plasticity. Adult neural plasticity has been showed in Lepidoptera and other insects (78–80), and we recovered several genes linked to axon regeneration (*APOD*), central complex development (*Ten-a, TENM2, DST, ALDH3A2, OGT*), central nervous system development (*disco*, *RpL4*), and neuropeptide hormonal activity (*tk*). Several genes were specific to retinal ganglion cell axon guidance (*TENM2*) and mushroom body development (*DAAM2*), leading us to speculate that plasticity could occur through neural wiring and plasticity are shaping sensory adaptation. Many Lepidoptera have distinct phenotypic plasticity in wing patterns and coloration in different seasons (81), and there is some evidence that they also have seasonal plasticity in their behavior and foraging preferences (82). Research has found Lepidoptera can override their innate preferences and learning preferences for new visual and olfactory cues after eclosion (83). It is possible the diel-niche and circadian rhythms too have plasticity, there are reports of some moth species like *Hyles lineata* showing relatively labile diel-niches possibly driven by temperature and resources (84–86). These same mechanisms allowing flexibility within a species, might be involved in diel-niche evolution between species.

### Circadian and behavioral regulators

The behavioral state of an animal to engage in any activity, e.g., feeding, flight or mating, is likely a function of circadian and behavioral regulators that respond to certain stimuli. We found differential expression of genes involved in locomotion (*unc-22, KCTD15,Tk*) and circadian or rhythmic behavior (*SREBF1/SREBF2, OGT, disco, JAK2*) in both species. We also found several key circadian regulators like *per*, *tim* although they were downregulated only in *Dryocampa*. C*lock-like* or *takeout* another gene under circadian control, (87) was expressed in both species, although without any significant up or down regulation. *Takeout* was moderately conserved in the moths we examined, but had diverged considerably among other insects, to the point where orthology searches with Orthofinder failed to recover orthologs among insects. Even among *D. melanogaster*, its closest homologs were *Jhbp5* and *takeout* which had 21-25% sequence identity (Supplementary Data 10,14). Surprising, it’s 3D structure was highly conserved (RMSD (align, super) = 2.556, 2.755, MatchAlign score (align, super) =199, 642.937, Supplementary Data 14, Fig. S12), highlighting an interesting example of likely functional convergence despite primary sequence divergence.

### Energy use and general cellular maintenance

In addition to the genes mentioned above, several key divergently expressed genes were involved in energy utilization. This finding makes biological sense, as once activity is initiated, energy mobilization and upregulation of basic cellular processes need to be maintained. Two of these genes were *SLC175A/MFS10* and *SLC2A6/Tret1-1*, which encode different trehalose sugar transporters.

Trehalose is an important sugar present in insect hemolymph (88). Other genes found were ribosomal (*RpL4, mRpS5*), golgi (*GCC*), and mitochondrial maintenance (*PARL/rho7*) along with transcriptional regulation (*Spt5)*.

### Disco as putative master diel-niche regulator in adult moths

Many genes that we discovered likely play a critical role in maintaining sensory shifts, but in order to identify genes that might regulate upstream, we looked for genes that were: 1) expressed in both species, 2) showed coincident expression patterns with respect to diel-niche, and 3) played a role in sensory and neural functioning and circadian control. Only *disconnected* (*disco*) fit these criteria.

*Disco* is a key developmental and patterning gene, first discovered for its role in neural migration from the disconnected optic lobe mutant phenotype (89). This gene is now considered an appendage development gene, with a well characterized pupal role in antennal and leg distal patterning in *Drosophila melanogaster* (*60*). Mutants also have a disrupted circadian locomotor rhythm due to the improper formation of neurons that express clock genes (90, 91). As a gene that is involved both in optic lobe, antennal formation, and in neural wiring and circadian control, it is a very strong candidate for driving behavioral diel-niche shifts and sensory adaptations (56, 57, 92). *Disco* expression data from *B. mori* suggests strong adult head, antennal, and nucleus expression, suggesting that it still acts as a transcription factor, and can regulate other genes (40).

*Disco* encodes the acid zinc-finger transcription factor and is approximately 500 and 1000 amino acids in length in *Drosophila* and *B. mori,* respectively. Our modeling showed that a ∼150 amino acid region is conserved structurally at the sequence level, and this region also contains zinc-finger motifs associated with DNA binding (93). This indicates that *disco* likely has retained its DNA binding and likely pupal and appendage patterning function in moths. We predicted the functional regions of *disco* in *Bombyx* based on evolutionary conservation modelling, and found an additional 500 amino acids, absent in *Drosophila*, were predicted to be functional (conserved and exposed). Domain and family level modeling predicted at least two additional zinc-finger domains in this region (Fig. S10,Fig. S11). We also found many phosphorylated sites surrounding the zinc-finger domains. Examining mutations between *Anisota* and *Dryocampa* revealed three mutations mapped to these predicted functional regions (Fig. 6B).

Given *disco’s* adult diel-specific expression in moths, additional zinc-finger DNA binding domains, and the high number of phosphorylated sites, we propose it as a candidate transcriptional regulator for diel-regulation in adult moths.

### Study species and their evolution and variation in life histories

*Anisota* and *Dryocampa* are closely related and ecologically similar saturniids of the subfamily Ceratocampinae, thought to have diverged ∼3.81 Mya (47). They largely occupy the same range and forest habitat in the Americas, feeding on large trees (*Anisota* on oaks and *Dryocampa* on maples) where population flares cause host plant defoliation (38). Because they reportedly hybridize (94), and pre- and postzygotic barriers are potentially ineffective, temporal partitioning may be important in their evolutionary history.

## Conclusion

This study provides a framework for how diel-niche evolution in Lepidoptera can occur.

Sensory and neural developmental genes appear to be key. We identify the pivotal role of the transcription factor *disco*, uncovering a second functional region with species-specific mutations in moths. Our findings provide useful targets for further genetic manipulation and highlight the insight non-model systems provide to genetics and development.

## Supplementary information

Supplementary tables and data are available for the reviewers and will be uploaded to repository [xxxxx] on submission.

## Author contributions

Y.S.: conceptualization, data curation, formal analysis, investigation, methodology, project administration, software, visualization, writing–original draft. R.M: investigation, methodology, project administration, data curation, writing - review & editing. A.J.: conceptualization, formal analysis, writing - review & editing C.S.: conceptualization, data curation, methodology, project administration, supervision. S.C.: formal analysis, software. K.G.: formal analysis, software, visualization, writing-review and editing, D.G.: investigation, project administration, data curation. R.L.: conceptualization, data curation, investigation, methodology, writing-original draft, writing -review & editing. C.H.: conceptualization, data curation, investigation, methodology, project administration, writing - review and editing I.K.: conceptualization, funding acquisition, methodology, resources, writing–review and editing. C.E.: formal analysis, software, visualization, data curation, writing- original draft, writing- review & editing. C.B.: software, validation, writing -review and editing. S.B.: funding acquisition, methodology, validation, writing–review and editing. J.T: conceptualization, funding acquisition, supervision, validation, writing–review & editing. A.K: conceptualization, methodology, resources, writing–review and editing, visualization, supervision, funding acquisition

## Supporting information

Supplemental Index 1

## Acknowledgements

We thank Jesse Breinholt, Martijn Timmermans and Andreas Zwick for help with project conceptualization. Kelly Dexter, Amanda Markee, and members of the McGuire Center for Lepidoptera and Biodiversity at the University of Florida assisted with animal care and wet-lab troubleshooting. We thank Belinda Pinto, Danielle DeLeo, Heather Bracken-Grissom, Jorge Perez-Moreno, Zhou Lei for discussions, advice and assistance with bioinformatics pipelines and wet-lab protocols. We thank Nicolas Alexandre, Sachit Daniel, Riddhi Deshmukh, David Plotkin, Chelsea Skojec and Nitin Ravikanthachari for comments and feedback on the manuscript. Janelle Nunez-Castilla, Jessica Liberles, Raghavan Venket, Yi-Ming Weng, Yu Fahong and Xiaokang Zhang assisted with troubleshooting bioinformatic pipelines and provided analytical recommendations. The authors acknowledge the University of Florida Research Computing for providing computational resources and support that have contributed to the research results reported in this publication (http://researchcomputing.ufl.edu). We acknowledge NBAF and UF-ICBR for assistance with sequencing and generating libraries.

Funding for the project was through NSF DEB-1557007, NSF PRFB-1612862, and NSF IOS- 1750833 and NERC NE/P003915/1. The Florida International University Presidential fellowship supported YS.

## Methods

### Insect rearing

Moths were reared under natural light-dark cycles at room temperature (25°C) and natural light conditions at the McGuire Center for Lepidoptera and Biodiversity (MGCL). Adults were sampled two days after eclosion at the two time points by C.H. and R.S.L.

### Sampling design

We collected 3-5 replicates (i.e., individual moths) per experimental treatment. We sampled at two time points per species at two midpoints of their circadian cycle at noon (‘day’) and midnight (‘night’) (Fig. 1, Table 1). Moths were allowed to eclose naturally and sexed, To ensure vision genes were not artificially stimulated, for collection at night we used red light, which is thought to not stimulate most Lepidoptera insect visual systems (95). We used a sterilized scalpel to remove the head, which was placed in a 1.5 ml Eppendorf tube, which was immediately placed in liquid nitrogen and stored at -80°C. It is worth noting that only males of most *Anisota* species are diurnal, including *A. pellucida*, our focal taxon. Females of all *Anisota* and both sexes of at least *A. stigma*, are nocturnal. In *Dryocampa*, both males and females are nocturnal. We therefore limited our transcriptomic work to males of *A. pellucida* and *D. rubicunda*.

### RNA extraction and quality control

Tissue was removed from liquid nitrogen and placed into a flat-bottomed cryo-tube with Trizol and 2-4 sterile, stainless-steel beads. The tissue was homogenized using a bead beater for 2.5 minutes at 1900 RPM. Beads were removed with a magnet, and chloroform added to separate phases.

Isopropanol precipitated the RNA, followed by several ethanol wash steps to pellet the RNA. Quality Control (QC) was undertaken with a high sensitivity Qubit assay. Samples with yields > 20 ng/μL were cleaned using a Thermo Scientific RapidOut DNA Removal Kit (Waltham, Massachusetts, USA). Sample quality was verified using Agilent 2200 Tapestation. Since heads comprised of a relatively small amount of tissue, initial extractions using kit-based methods resulted in low yields.

Several rounds of troubleshooting were required before completion of extraction using a modified Trizol based protocol with a DNA clean up step. We measured the quantity of RNA using a Qubit fluorometer and used Agilent tapestation or Bioanalyzer to characterize extraction quality, weight, and size distributions. Where possible, a concentration of more than 10 ng/μL was used for library preparation. Yields below 10 ng/μL were prone to higher loss when purifying with a DNAse agent, which often caused 20-30 % loss in RNA yield, approaching 5ng/μL is below the Qubit detection threshold.

### Library preparation and NGS sequencing

Libraries were prepared by sequencing cores at NBAF-Liverpool using the NEB polyA selection and NEBNext Ultra II directional stranded kit, suitable for low input yields. 12 samples were run on one lane of an Illumina NovaSeq using SP chemistry (Paired-end, 2x150 bp sequencing). Samples were shipped overnight from MCGL to the NERC-NBAF after dehydrating in a biosafety chamber using GenTegra RNA tubes.

### Read trimming and cleanup

For *Anisota pellucida* and *Dryocampa rubicunda*, trimming was undertaken by the NERC-NBAF core and the raw Fastq files were trimmed for the presence of Illumina adapter sequences using Cutadapt version 1.2 (96). The option -O 3 was used, so the 3’ end of any reads that match the adapter sequence for 3 bp or more were trimmed. Trimming was also done with trimmomatic. The reads are further trimmed using Sickle version 1.200 with a minimum window quality score of 20. Reads shorter than 15 bp after trimming were removed.

### Transcriptome library sizes, QC, and PCA

Quality Control (QC) was conducted on the trimmed reads, library size varied for each species (Fig. S1). We examined expression data and removed genes with TPM < 1. PCA results showed that *Anisota* and *Dryocampa* did not separate with diel-niche (Fig. S3).

### *De novo* assemblies

We combined reads from multiple samples and generated several reference de novo transcriptomes using different assemblers, and then combining them, compared their quality using BUSCO scores and number of single-copy BUSCO genes across the different versions (see Table 2 for the BUSCO scores, redundancy and duplication for the various assemblies). We found that unfiltered assemblies were highly redundant, but had the highest BUSCO scores. These assemblies were too overrepresented to be used in any downstream analysis, so we used Transrate, CD-HIT, MMeqs2, as well as Transdecoder, to filter the assemblies. We chose v5 and v6 to use in downstream analysis, listed as the last two assemblies for each species (Table 3) and repeated certain analyses with both assemblies to see how stringent or a less stringent filter would affect the results.

### Multi assembler *de novo* transcriptome assembly

To our knowledge, there is no publicly available annotated reference genome for the two species used in this study at the time the analyses were performed; therefore, we chose to build a de novo transcriptome assembly with the pre-processed reads using the NCGAS pipeline, which combines multiple assemblers and then uses an evidence-based gene modeler to decide on the final set. For each species, we combined forward and reverse reads from all samples and normalized them using a normalization script in Trinity v2.12.0 (97). Normalized reads were used as input to generate assemblies using Velvet v1.2.10 (98), TransAbyss v2.0.1 (99), SOAPdenovo v1.03 (100) with various kmer sizes. These assemblies were combined and filtered using Evigenes v4: 2020, March (101, 102), transdecoder and filtered with scripts provided in the NCGAS pipeline (103).

### Transcriptome QC, annotation and filtering

Further quality control and filtering was conducted using BUSCO (104), transrate (105), QUAST (106) (v5.02), CD-HIT(v4.6.8) (107, 108)and cascaded clustering with MMseqs2 (v12) (109). BUSCO scores were greater than 95%, but many genes had multiple versions from the different assemblers, and because this would bias the downstream analyses we attempted to reduce the redundancy at the cost of lowering BUSCO scores. Protein coding regions were predicted using TransDecoder (https://github.com/TransDecoder/TransDecoder) and the cds file from was run through Transrate (105), which further pulled contigs based on mapping rate and identified a “good” collection of transcripts, the resulting protein-coding sequences were filtered with additional CD-HIT (107, 108)and MMseqs2 (109) filtering. This resulted in lower BUSCO scores 75-85% but also much lower redundancy in the gene set of about 1-3 fold. We ran the WGCNA (52) and GO enrichment (110) analyses on the more stringent assemblies (BUSCO 75%, 2% genes with duplication) since they are more sensitive to redundancy, but RNA-seq was run across the slightly lower stringency analyses to be more inclusive (BUSCO∼85%, 60-70% genes with duplications).

### Annotation of reference transcriptomes

Reference genomes were annotated with eggNOGmapper and Orthofinder (54, 55). We also tried annotating the reference transcriptomes with Trinotate (111), which run against the swissprot and pfam databases using hmmscan, blastx, blastp, signalP and tmmhmm, KEGG, and GO and eggNOG (49, 110, 112–120). ORF predictions for orthofinder were obtained using *Antheraea pernyi* as a model. Orthofinder was run with well-annotated Bombycidae and Saturniidae moths, *Bombyx mori*, *Antheraea pernyi*, and *Antheraea yamamai*, to generate clusters. *Bombyx mori* was a useful reference since the Orthofinder cluster for each species generated a list of mostly single-copy genes for which more Lepidoptera annotations were available from SilkDBv3 (40). While this was not as complete as trinotate, the annotations provided from this procedure were more accurate than Trinotate. Trinotate annotation had many more human/ vertebrate hits than insect matches than eggNOGmapper and Orthofinder transferred annotations, so while the files are provided as a reference, they were not used in downstream analyses.

### Differential gene expression (DGE)

Reads were mapped using Salmon (121) and differential gene expression analysis was conducted using EdgeR (122) and DESeq2 (123). EdgeR(v 3.38.1) was used to normalize and test for significantly expressed genes. DESeq2(v1.36.0) was also used to normalize and to generate PCAs and other summary statistics. We used ’Day’ as the treatment and ’Night’ as the control for all groups with EdgeR, although the order of fold change switched for DESeq2. The p-values obtained were adjusted to account for multiple hypothesis testing with FDR for EdgeR and adjusted p-value from DESeq2. Genes with FDR or adjusted p-values <0.05 and |FC| > 2 were used as a criteria to identify significantly expressed genes.

### Gene enrichment analysis

We performed gene enrichment analysis using GO terms with the tools TopGO v2.48.0 (124) and ShinyGo v0.75c (125) (http://bioinformatics.sdstate.edu/goc/). This analysis involved comparing the selected genes of interest to a gene universe with GO term annotations to determine if there were overrepresented or underrepresented GO terms. To obtain genes with GO annotations, we utilized similarity clusters to map annotations from *Bombyx mori*.

In TopGO, we employed a more stringent approach by using filtered transcriptomes to mitigate bias from gene duplications. We selected genes of interest based on a False Discovery Rate (FDR) threshold of less than 0.05. We ran two different enrichment algorithms, ’classic’ and ’elim’, and used two test statistics, Fisher’s exact test and a Kolmogorov-Smirnov-like test. The tests were performed for both ’Biological Process’ (BP) and ’Molecular Function’ (MF) categories. While we generated tables of significantly enriched GO terms, obtaining individual genes within each group was limited due to the constraints of TopGO with custom annotations.

For ShinyGO, we utilized the custom v0.75c which included the updated *Bombyx mori* genome stringent. For all analysis, we used the following settings (pathway size: min.=1, max=2000). The settings for number of pathways to show and FDR cut-offs were chosen to represent the entire list of top 100 genes in the final datasets, although in some cases a fewer number are represented in the visualizations. We focused on the ’Biological Process’ (BP) category for the various tests. Tables of GO terms, gene information, and graphs summarizing the significantly enriched GO terms were generated. We repeated several different analyses **1)** A less stringent analysis was run where genes with FC<|2| and padj. <0.05. This was useful to visualize the genes up and downregulated functionally and the background used was all the genes recovered from the DEA analysis that had orthologs. An FDR cut-off <0.1-0.3. For the background gene set, we used the species’ de-novo transcriptome matching *Bombyx* ortholog set.

**2)** A more stringent criteria was used to examine highly upregulated genes a FC<|5| and padj. <0.05 cutoff was used, and filtering only genes that occurred in both EdgeR and DESeq2 analyses. We selected a single representative when multiple Bombyx orthologs were recovered, using the ortholog with the most complete annotation. An FDR cut-off <0.17 was used. For the background gene set, we used the species’ de-novo transcriptome matching Bombyx ortholog set.

**3)** For WGCNA clusters tan and grey60, we used their respective species background expressed transcript set with a FDR cutoff <= 0.1. For blue and turquoise, the background of all orthologs found in *Bombyx mori* was used. An FDR cutoff<1 was used, since smaller values provided insufficient genes given the larger number of genes being tested.

### Gene network analysis

Gene network analysis was undertaken with WGCNA (1.71) (52, 126). DeESeq2 was used to normalize reads and data was formatted with tidyverse. WGCNA identifies modules of co-expressed genes. We chose modules that showed clear day-night differences and mapped the GO terms with Revigo and ShinyGo. We also combined counts from both species and tested for clusters. Here we chose modules that showed species specific clusters.

### Gene annotation and filtering

DEGs, enriched genes and or co-expressed transcripts/ genes were annotated by cross referencing them with the *B. mori* annotations (see above). Since *B. mori* mappings were not always ’one-to-one’, mappings that were ’one-to-many’ were dropped, i.e., transcripts that mapped to multiple *Bombyx* genes were dropped. We also improved annotation by mapping the overlapping transcripts with eggnog mapper. We further filtered these genes into sub functional categories, by identifying genes with GO annotations or descriptions linked to vision, smell, hearing, circadian, behavior and brain the list of go terms for each group was obtained by querying amigo (http://amigo.geneontology.org/) (Table 8).

### Gene Mining and in-silico evolution

We mined genes of interest from ∼bombycoidea moths and closely related families that had well annotated genomes on Ensembl and NCBI (Supp Table 6) and across model insects (Supp. Table 7). These genomes were chosen as all Bombycoidea from Ensembl with well annotated assemblies and five species Noctuidae and Geometridae each were included as close relatives. We added *Bombyx mori* as a reference and *Antheraea pernyi* to represent Saturniidae, and their peptide files were taken from their respective source papers (40–42). For insects, we chose long-read well annotated genomes from Ensembl (Ensembl Gene build) and five model systems from insect base (127). We ran Orthofinder (v2.5.2) (49, 55) with ‘orthofinder -f bom-rel/ -S diamond -M msa -A mafft -T fasttree -a 1 -X -z -t 16’ to recover orthologs. Each orthogroup often contained multiple sequences per species. We chose the sequence with the highest identity to the reference *Bombyx mori* sequence using custom python scripts. In the rare instances where there were multiple references, we took the longest one. To model the 3D structure of each reference *Bombyx mori* sequence, we ran AlphaFold v2.1.2 (128) using Deepmind’s run_alphafold.py (https://github.com/deepmind/alphafold/blob/main/run_alphafold.py) on University of Florida’s HiperGator. Model database files (i.e. BFD, MAGnify, PDB70, mmCIF PDB, UniRef30, UniRef90) downloaded from Deepmind in February 2022 were used as parameter value inputs. All other parameters were used with default settings. The relaxed predicted model with highest ranked pLDDT (per-residue estimate corresponding to model’s predicted score on the lDDT-Cα metric) was chosen as the final model (129).

We calculated a conservation score for each alignment using Alistat. For alignments less than 1500 amino acids and with high conservation score (Proteins larger than 1300 aa were difficult to model without high memory GPU’s), we modeled conservation using Consurf (130–134). The results were displayed using Jmol first glance viewer (http://firstglance.jmol.org/) and as screenshots from the default viewer. The output for alpha fold runs and consurf is available in the (Supplementary Data). PyMol (135) was used to align the 3D structures and Aliview was used to compare alignments. We used InterProscan (https://www.ebi.ac.uk/interpro/about/interproscan/) and NetPhos3.1 (136, 137) for predicting the domains and phosphorylated sites.

