## Supplemental Index 1 for "The developmental gene *disco* regulates diel-niche evolution in adult moths"

**This PDF file includes:**

Supporting text
Figures S1 to S12
Tables S1 to S8
Legends for Datasets S1 to S14

**Other supporting materials for this manuscript include the following:**

Datasets S1 to S12: Available at
https://www.dropbox.com/sh/fhh9fwcgnlnwth2/AAACRtDvd52_A5qGiqWOzSZma?dl=0

### Supplemental Figures

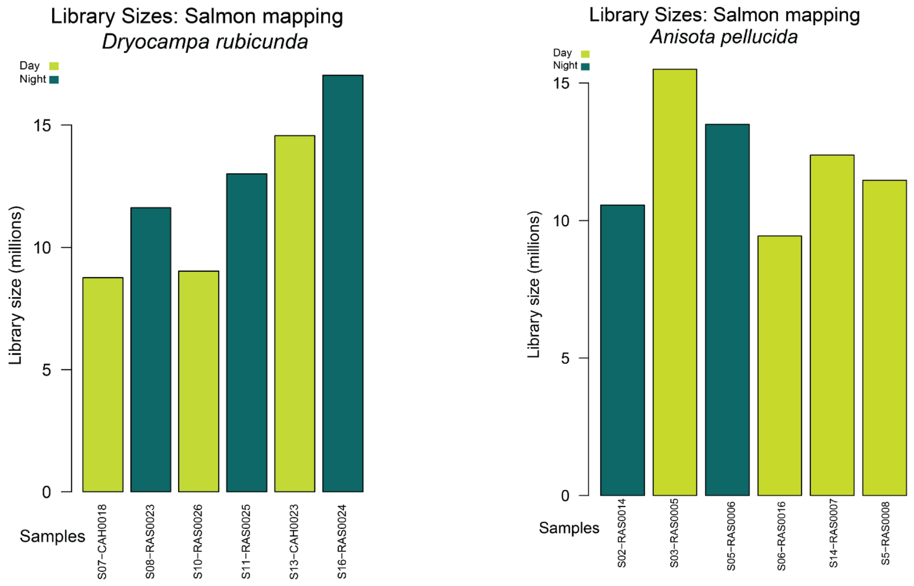


**Figure S1**: Variation in salmon mapped libraries for *Dryocampa rubicunda* and *Anisota pellucida*

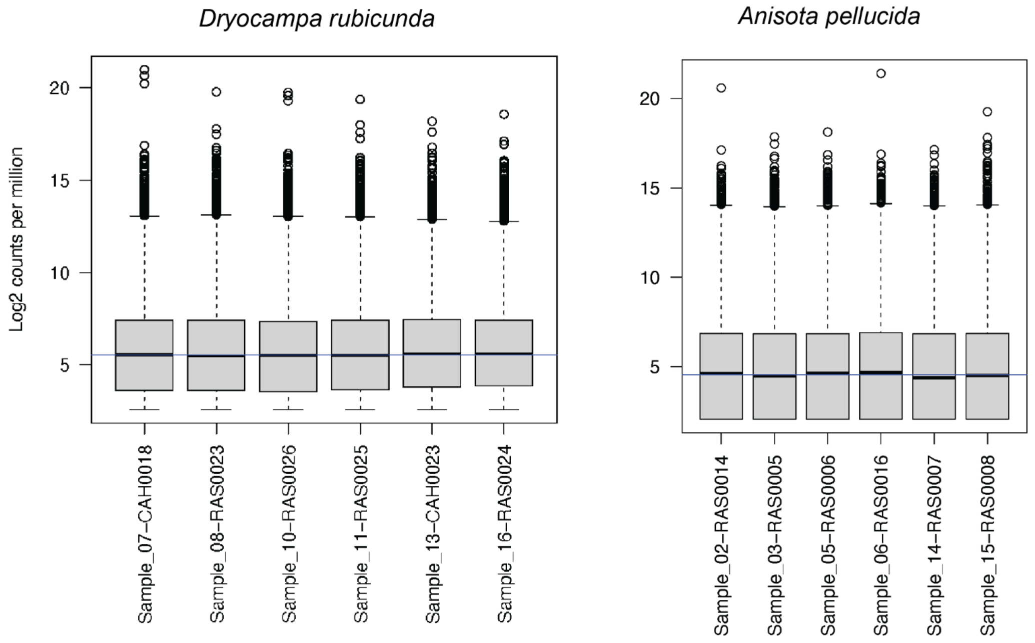


**Figure S2**: Variation in normalized libraries for *Dryocampa rubicunda* and *Anisota pellucida*

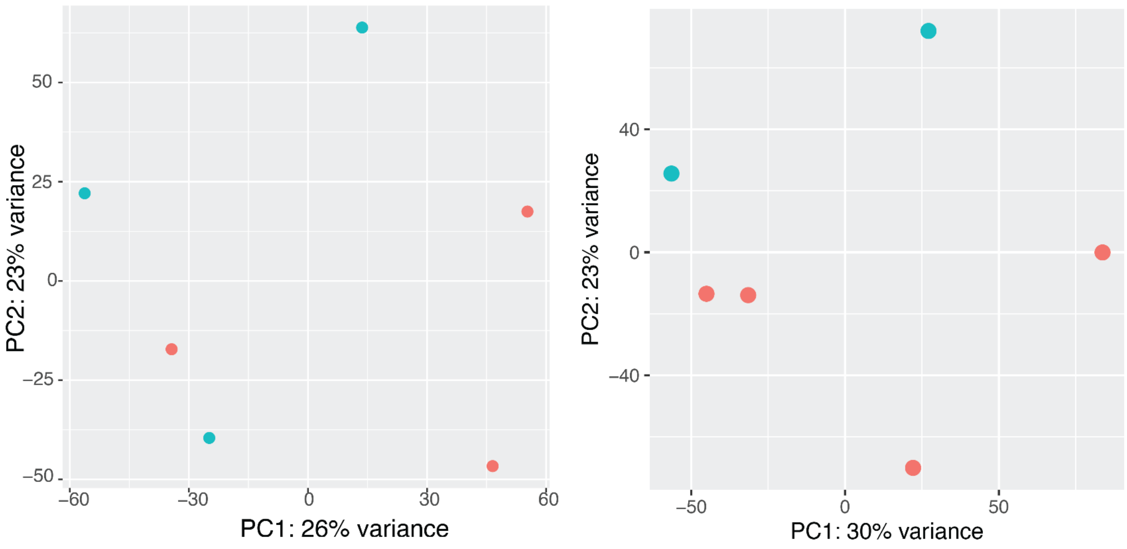


*Dryocampa rubicunda* *Anisota pellucida*

**Figure S3**: PC1 and PC2 of libraries labelled for *Dryocampa rubicunda* and *Anisota pellucida*. Orange and blue labels refer to midday ‘Day’ and midnight ‘Night’ collection time points, respectively. This PCA was generated from mapped salmon reads after normalising with DESeq2.

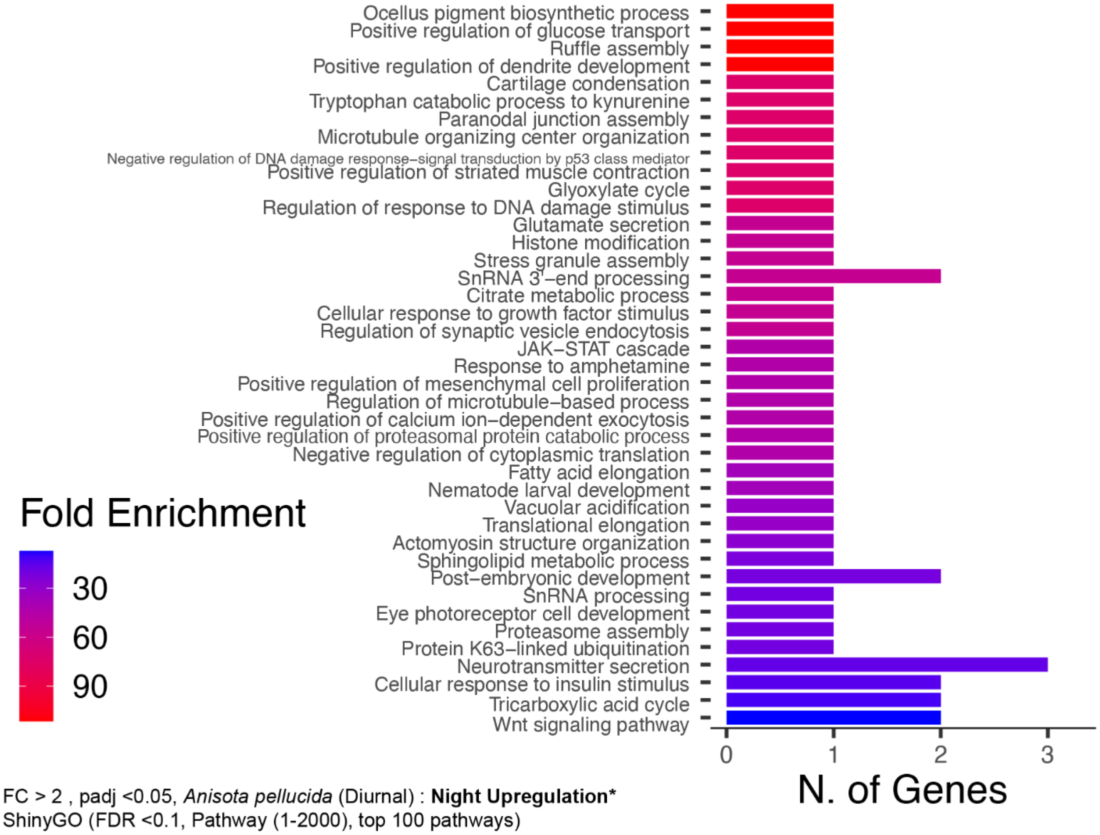


FC > 2 , padj <0.05, *Anisota pellucida* (Diurnal) : **Night Upregulation***
ShinyGO (FDR <0.1, Pathway (1-2000), top 100 pathways)

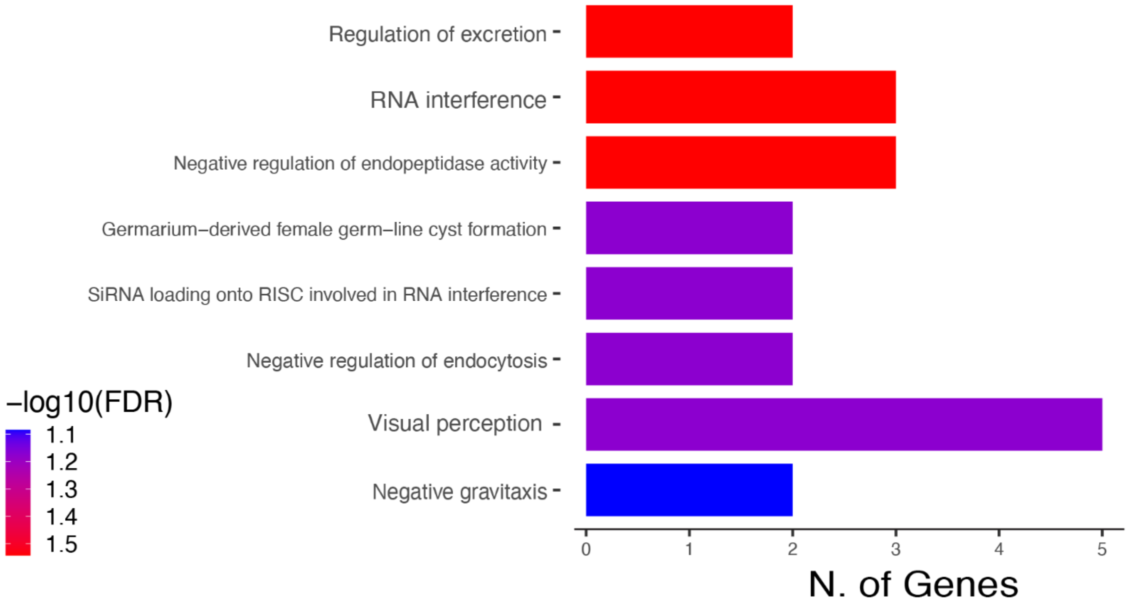


FC < -2 , padj <0.05, *Anisota pellucida* (Diurnal) : **Day Upregulation***
ShinyGO (FDR <0.1, Pathway (1-2000), top 20 pathways)

**Figure S4**: Enriched lists of GO terms associated with significant positively and negatively expressed genes recovered from 2 or more analyses (DESeq2, WGCNA, EdgeR) for *Anisota* *pellucida.* ShinyGo was used to highlight terms with genes showing representation compared to the background of detectable genes values representing ShinyGo overrepresentation. Top: Enriched Day Upregulated genes (FDR>0.1) with gene expression FC>=2, adjusted p-value <0.05. Bottom: Enriched Night Upregulated genes (FDR>0.1) with gene expression FC<=2, adjusted p-value <0.05 and that showed at least 2 occurrences in the combined dataset. Values are sorted by -log FDR *Genes that showed up in at least 2 separate analyses are included, (>=2 occurrences in the combined dataset). The background used was all transcripts that were recovered in the DEG analysis and had *Bombyx* orthologs

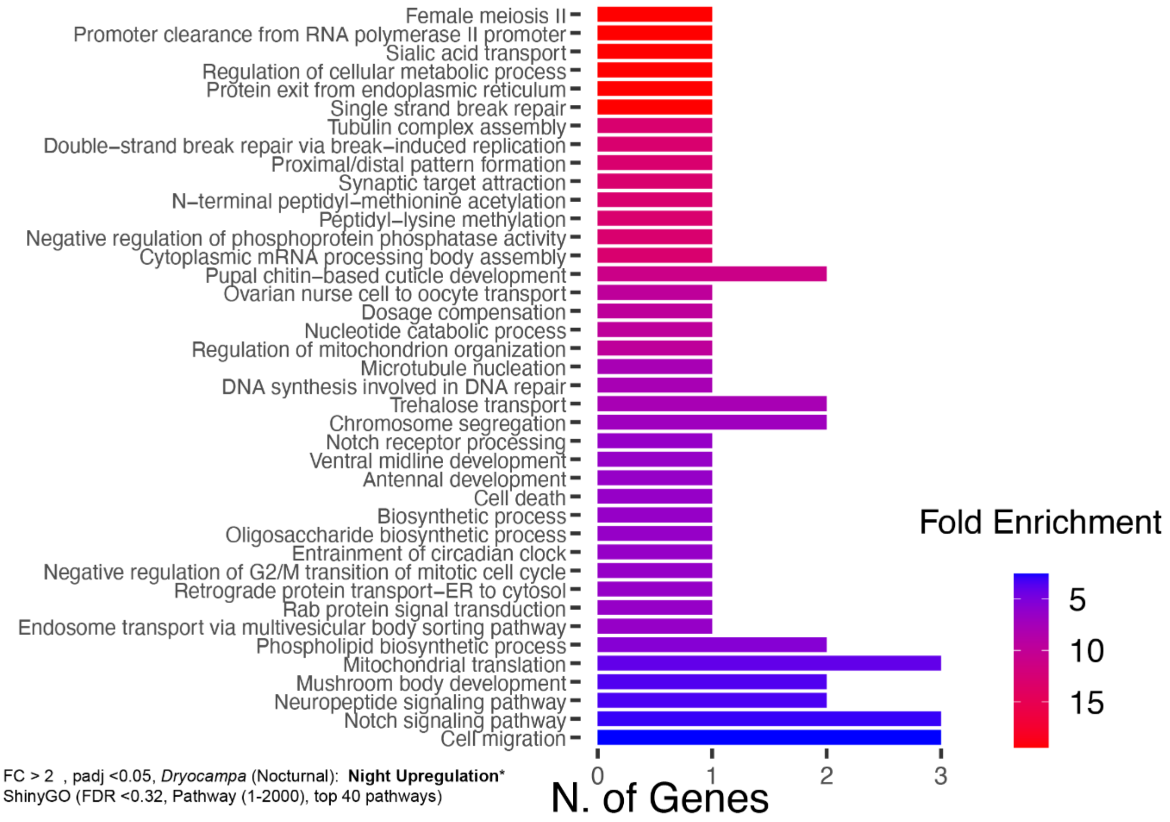


FC > 2 , padj <0.05, *Dryocampa* (Nocturnal): **Night Upregulation***
ShinyGO (FDR <0.32, Pathway (1-2000), top 40 pathways)

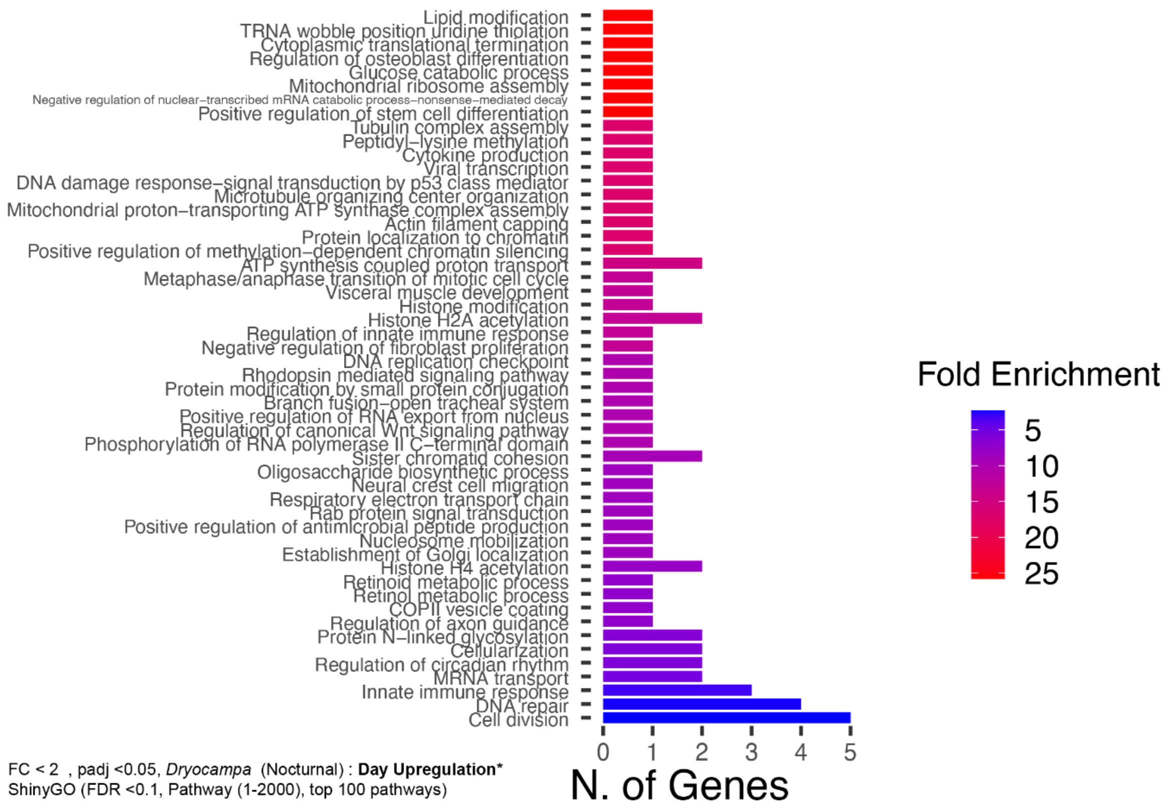


FC < 2 , padj <0.05, *Dryocampa* (Nocturnal) : **Day Upregulation***
ShinyGO (FDR <0.1, Pathway (1-2000), top 100 pathways)

**Figure S5** : Enriched lists of GO terms associated with significant positively and negatively expressed genes recovered from 2 or more analyses (DESeq2,WGCNA, EdgeR) for *Dryocampa rubicunda* . ShinyGo was used to highlight terms with genes showing representation compared to the background of detectable genes values representing ShinyGo overrepresentation. Top: Enriched Day Top 40 upregulated genes (FDR>0.32) with gene expression FC>=2, adjusted p-value <0.05, 77 terms enriched , but not all are displayed, see supplementary material for entire list. Bottom: Enriched Night Upregulated genes (FDR>0.3) with gene expression FC<=2, adjusted p-value <0.05 and that showed at least 2 occurrences in the combined dataset *Genes that showed up in at least 2 separate analyses are included, (>=2 occurrences in the combined dataset). The background used was all transcripts that were recovered in the DEG analysis and had *Bombyx* orthologs

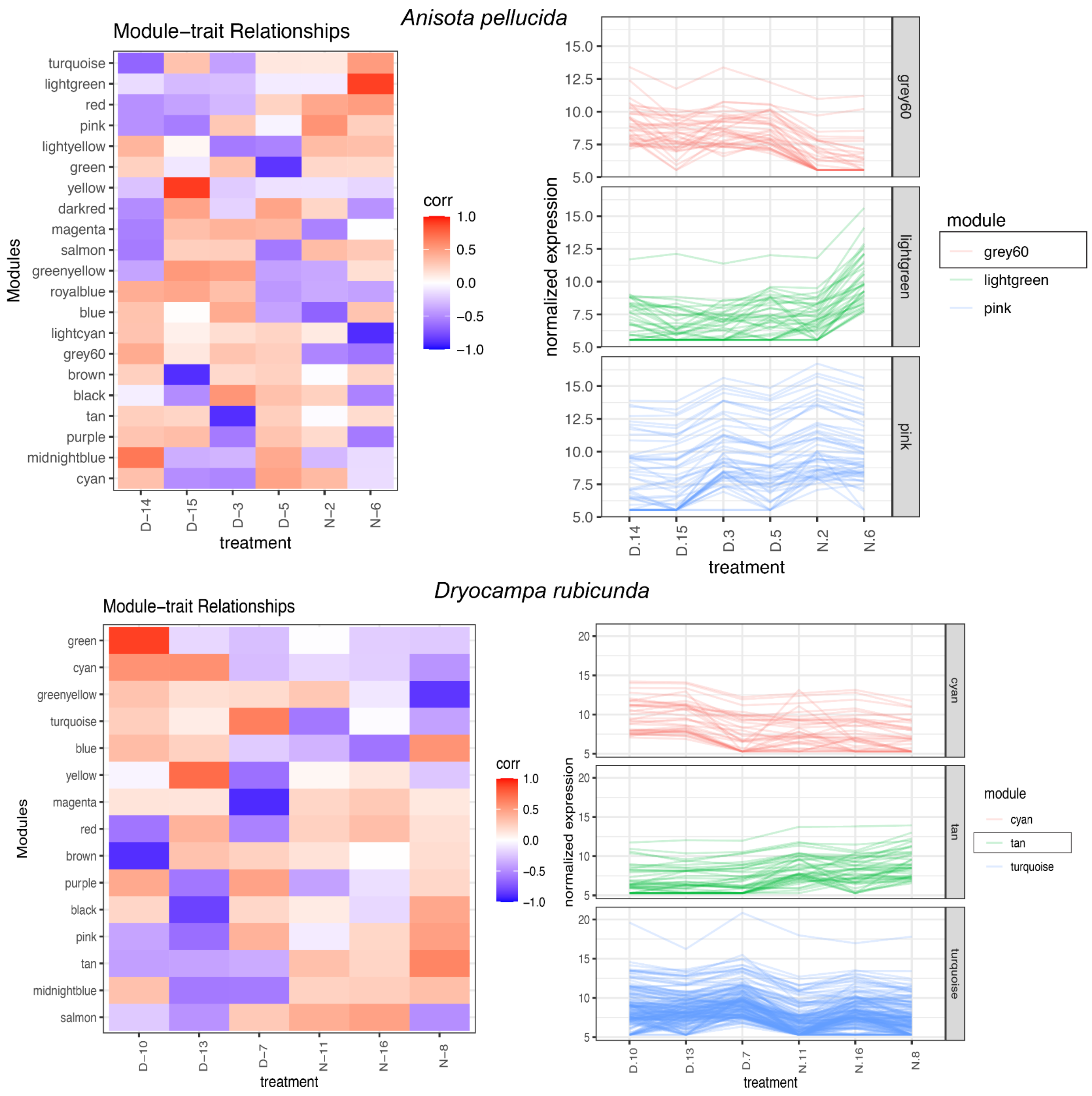


**Figure S6:** Modules of clustered genes grouped using normalized expression for *Anisota pellucida* (top) and *Dryocampa rubicunda* (bottom) Grey60 (38 genes) and Tan (32 genes) modules show species- time of day linked patterns. Left: Shows how patterns of gene expression correlate across samples for modules Right: Shows the normalized expression for all. Normalization was done with DESeq2 and reads were mapped to the more stringently filtered transcriptome. A soft power analysis was done and the picked power=9 for the WGCNA analysis.

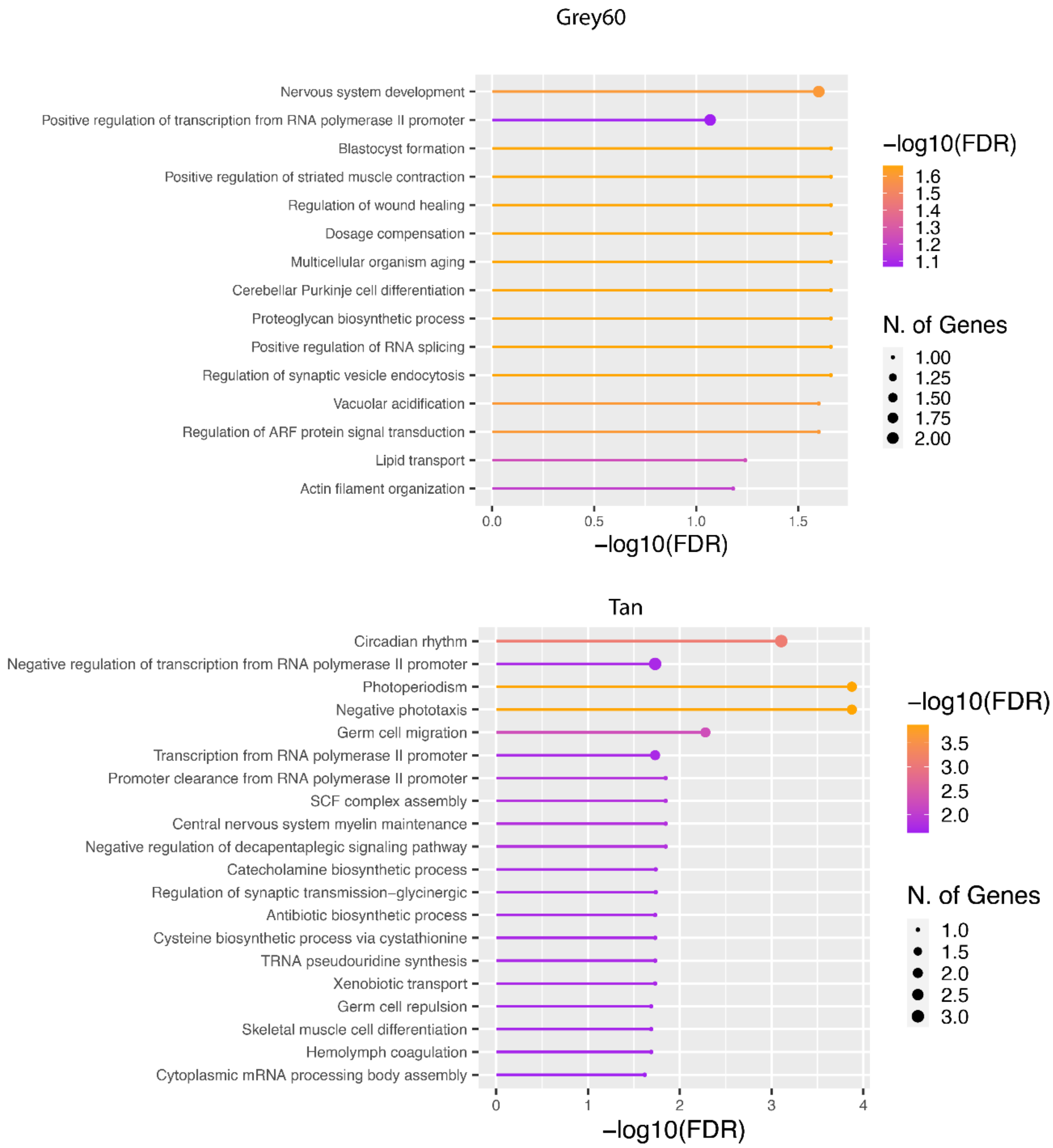


**Figure S7:** Shiny Go enrichment for Modules of clustered genes grouped using normalized expression for *Anisota pellucida* (top) and *Dryocampa rubicunda* (bottom) Grey60 and Tan. Background used was all *Bombyx mori* genes and FDR cut-off =0.1 min pathway =1 and GO Biological processes are displayed.

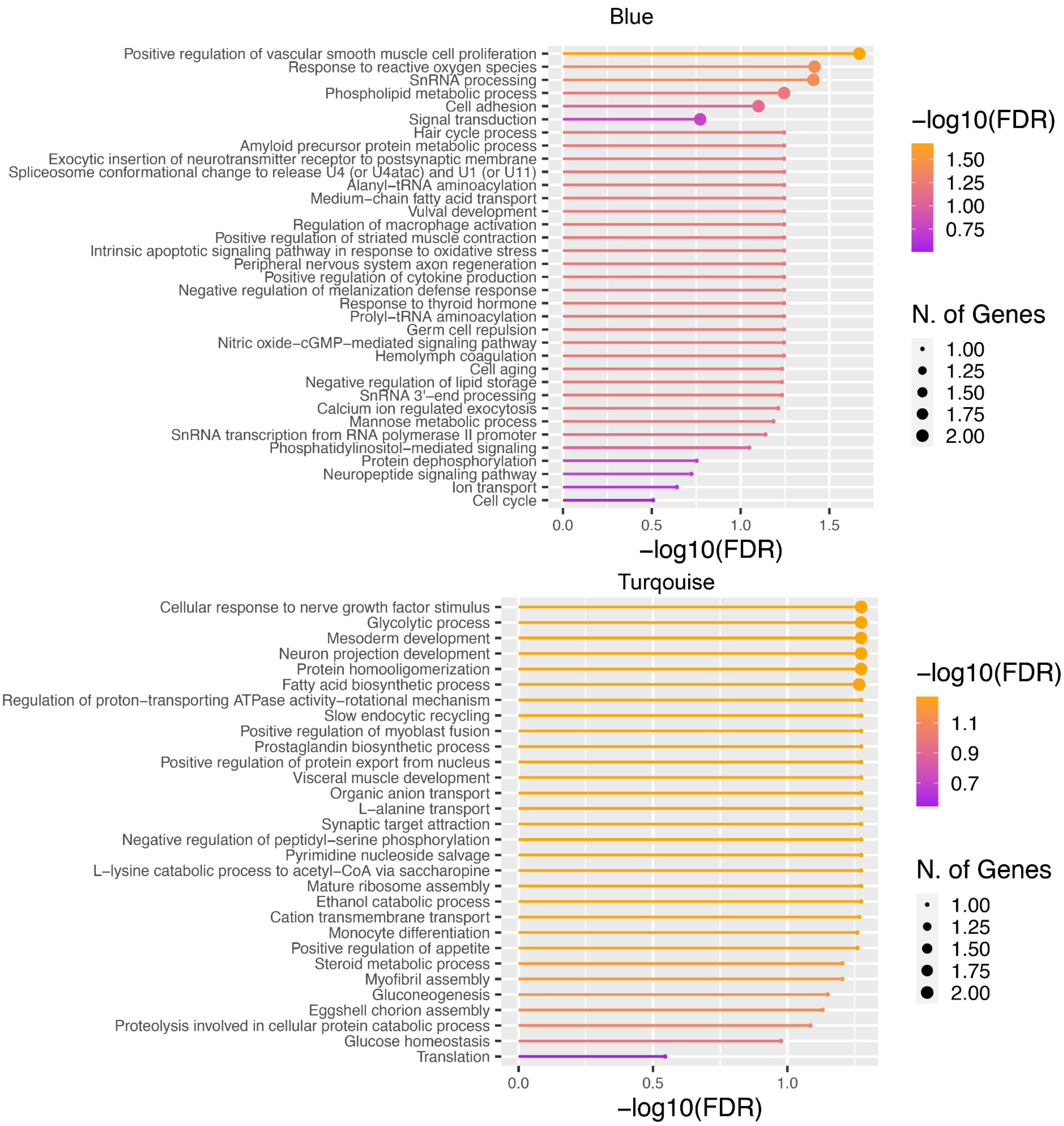


**Figure S8:** Shiny Go enrichment for Modules of clustered genes grouped using normalized expression for combined overlapping reads for the two species (Top) Blue and (Bottom) Turquoise. Background used was all *Bombyx mori* genes and FDR cut-off =1 min pathway =1 and GO Biological processes are displayed.

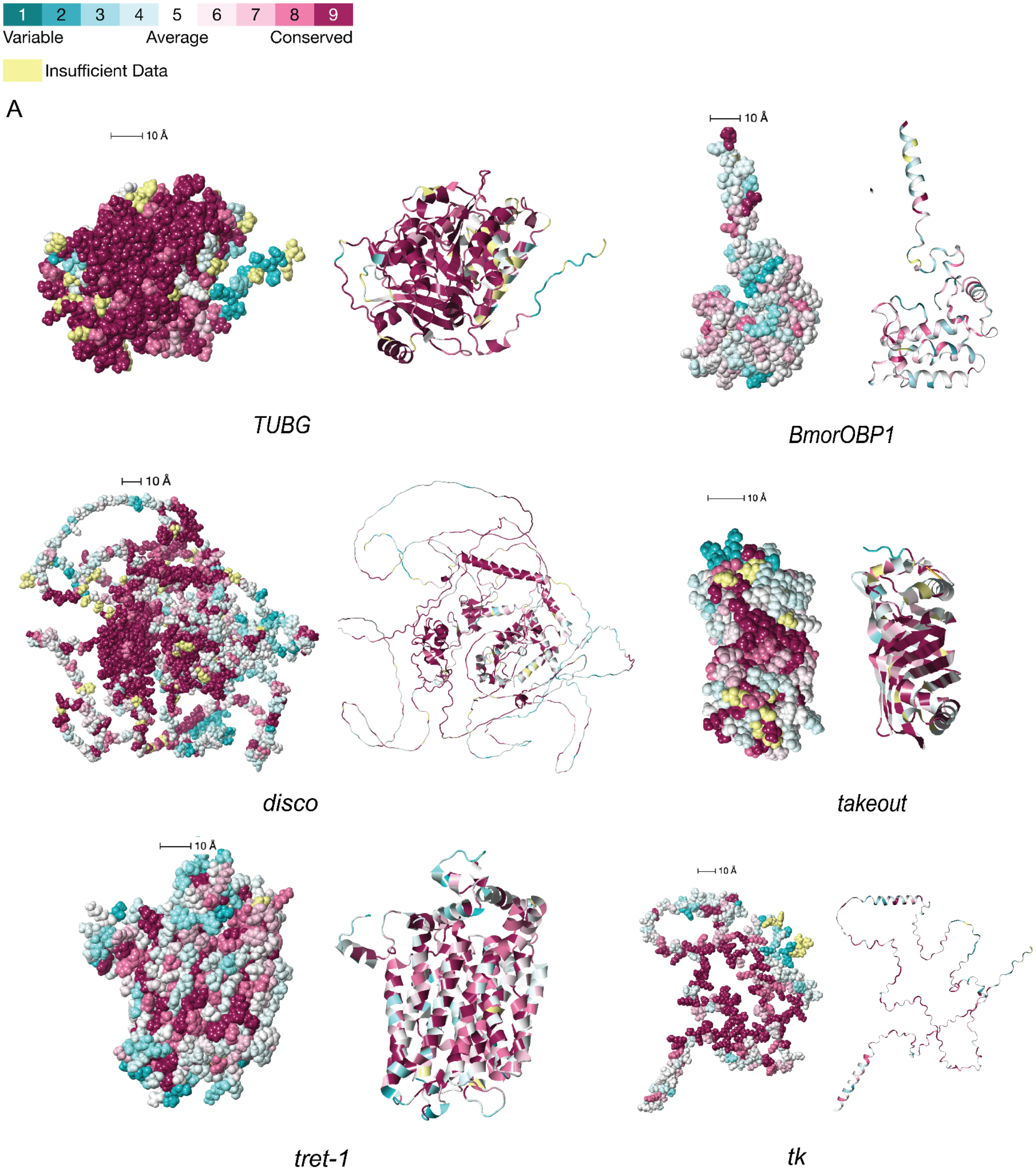


**Figure S9:** Variation in structural conservation across moths for predicted proteins for genes interest. Structures were obtained using AlphaFold predictions of *Bombyx mori* homologs. Left: Space filling models, Right cartoon models, colors represent conservation across the alignment from or Bombycoidea and related moth species. Conservation scores were calculated and mapped onto the alpha fold structures using the Consurf (131) web server (For other proteins, see Supplementary data)

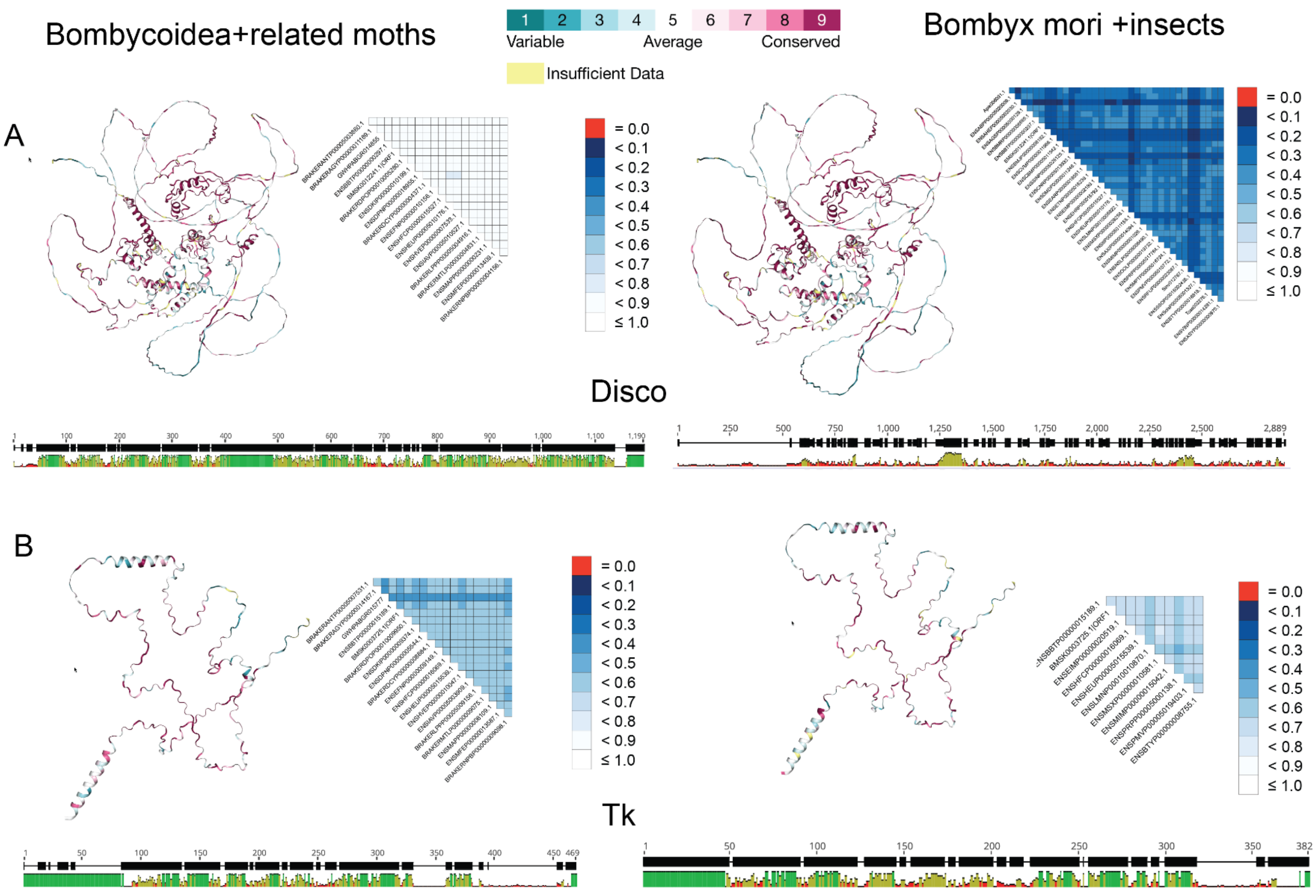


**Figure S10**: Illustration of conservation across the alignments and structure for **(A)** *disco* and **(B)** *tk* across Bombycoidea and relatives (Left) and insects (Right). Each inset contains the *Consurf* mapped pdb structures on the left. The *alistat* pairwise sequence conservation in the alignment scores on the right and the consensus of the protein alignments were visualized using *Geneious.* For conservation scores for other genes and lists of the species used for this modelling, see Supplementary Tales (Supp. Table: 5-7). Supplementary Dataset 10 has the associated models and output from the analyses.

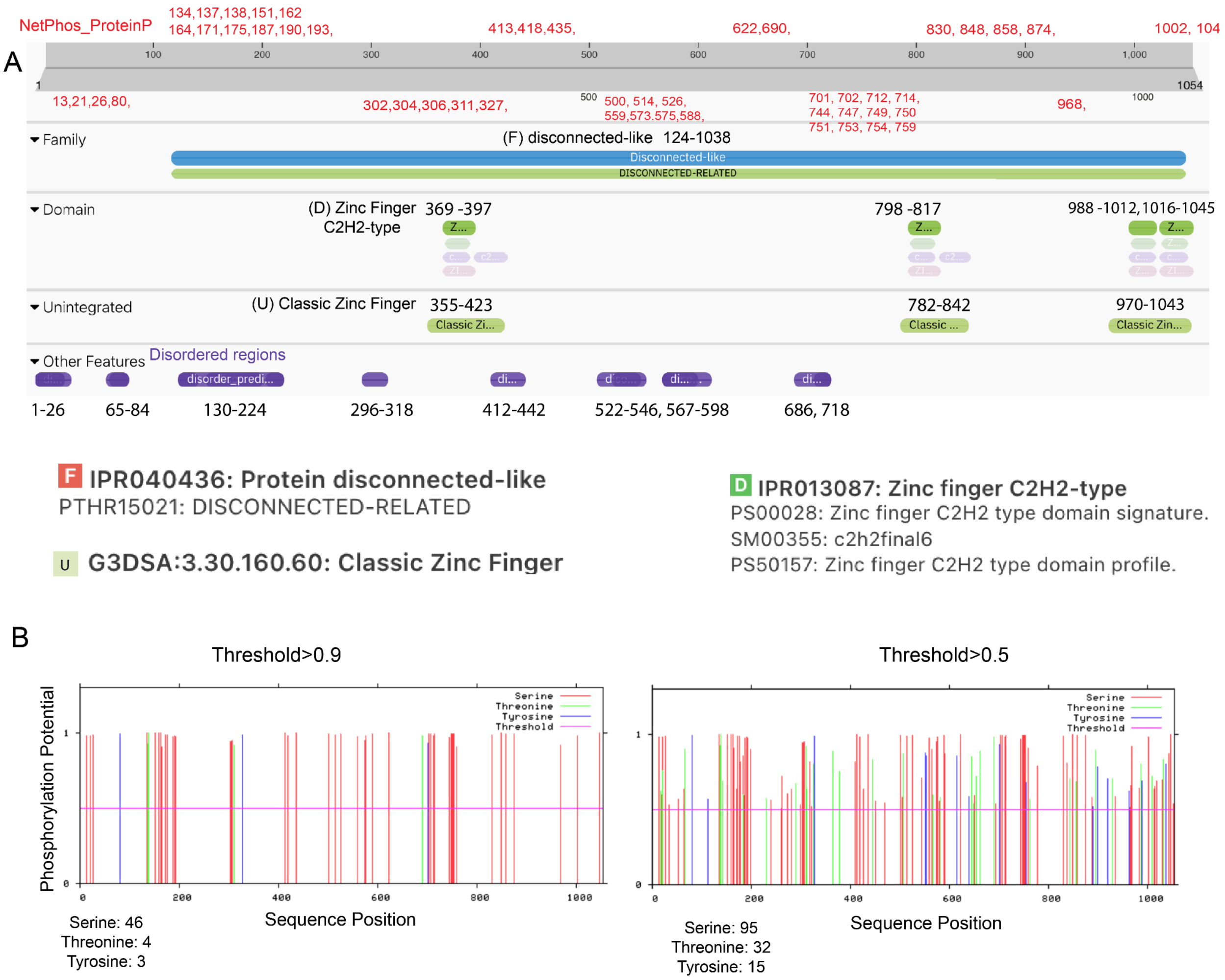


**Figure S11:** InterPro and NetPhos predictions for *B. mori disco* protein. **(A)**InterPro predicted several zinc-finger domains, disordered regions and disco and disco-related families. The numbers in black highlight the regions for the predicted domain and the numbers in red list the NetPhos predicted phosphorylated sites (threshold >0.9). **(B)** An overall distribution of the predicted phosphorylated sites with different thresholds (Left, Threshold=0.9, Right Threshold= 0.5)

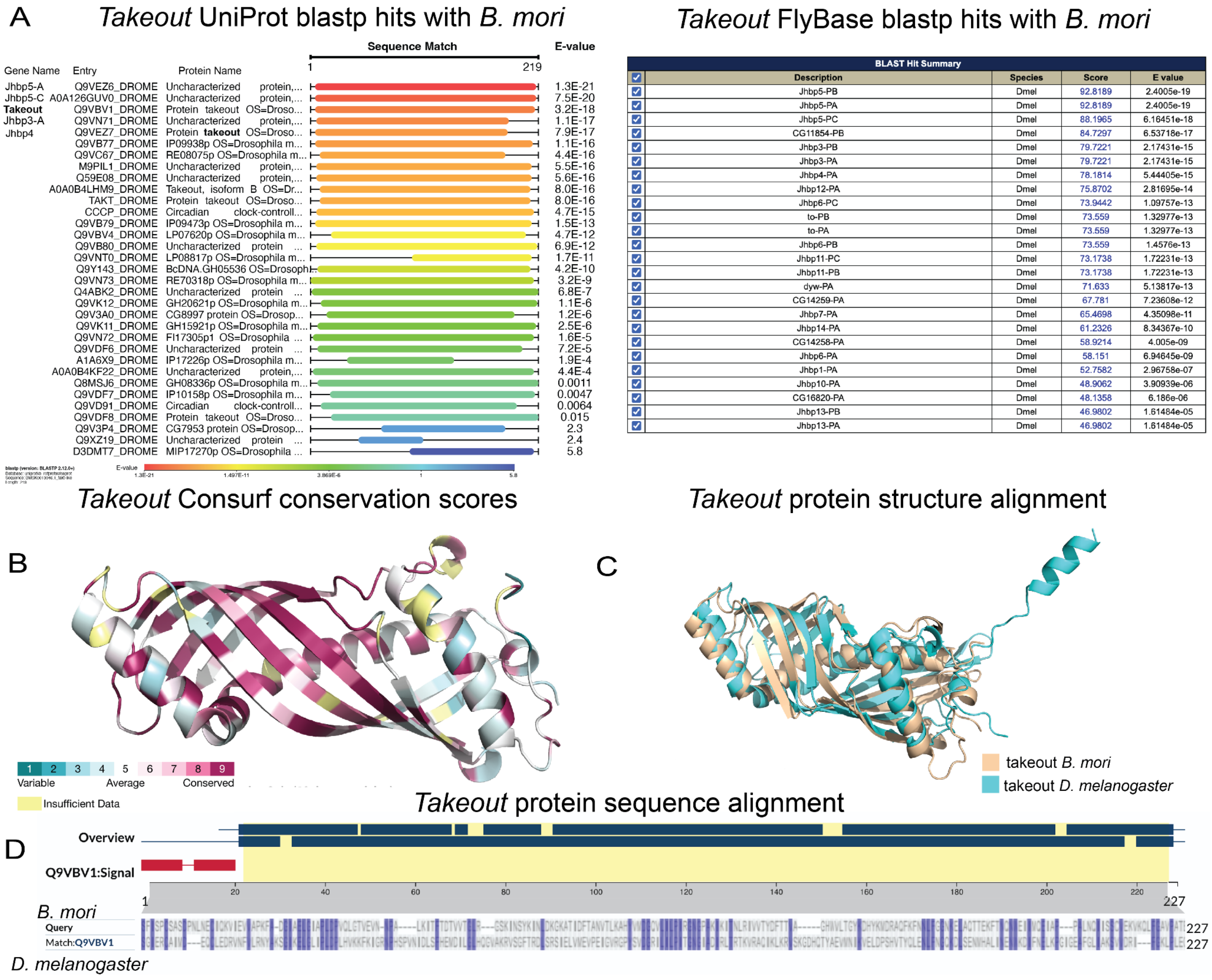


**Figure S12:** Lack of sequence level conservation in takeout, but high structural overlap **(A)** Right: Takeout UniProt blastp hits. (BLOSUM62, filtered for *D. melanogaster*), alignments =250, e-threshold= 10, database = uniprotkb_refprot+swissprot, list of illustrative hits is shown. Left: Similar blastp results from flybase limiting the species to *D. melanogaster*. **(B)** Consurf predicted scores for closely related moths mapped on to the *B, mori* takeout predicted protein structure. **(C)** 3D structural alignments for fly (*D. melanogaster)* and moth (*B. mori)* takeout 3D models. **(D)** Uniprot protein sequence alignment for fly and moth takeout proteins.

**Supplementary Tables**

| Species | Diel-niche | Collection Time | Replicates (n) |
|---|---|---|---|
| *Anisota pellucida* | Diurnal | Day | 4 |
| | | Night | 2 |
| *Dryocampa rubicunda* | Nocturnal | Day | 3 |
| | | Night | 3 |

**Table S1:** Sampling design showing species, collection time and number of replicates in each treatment. All specimens were male.

| Species | Assembler | kmer-size | Max- BUSCO groups | Lep BUSCO Score(%) | max-N50 |
|---|---|---|---|---|---|
| Ani_pel | Velvet | 35-85 | 4703 (k55) | 88 | 3918 (k55) |
| Ani_pel | Soap | 35-85 | 4682 (k35) | 88.5 | 1661 (k35) |
| Ani_pel | Transabyss | 35-85 | 4216 (k35) | 79 | 1332 (k35) |
| Ani_pel | Trinity | 25 | 4948 | 93 | 1608 |
| Dry_rub | Velvet | 45-85 | 4768 (k55) | 90 | 3957 (k55) |
| Dry_rub | Soap | 35-85 | 4753 (k35) | 89 | 1915 (k75) |
| Dry_rub | Transabyss | 55-85 | 3159 (k55) | 59 | 1058 (k55) |
| Dry_rub | Trinity | 25 | 4939 | 93 | 1595 |

**Table S2:** Assembly variation using different assemblers and kmer settings.
Key: Dry_rub: *Dryocampa rubicunda*, Ani_pel: *Anisota pellucida.* All models included day and night samples.

| Species | Method | Total DEGs | DEGs (Up) | DEGs (Down) | Annotated DEGs | Unannotated % |
|---|---|---|---|---|---|---|
| *A. pellucida D* | EdgeR | 350 | 154 | 196 | 211 | 39.71 |
| *D, rubicunda N* | EdgeR | 393 | 211 | 182 | 236 | 39.95 |
| *A. pellucida D* | DESeq2 | 697 | 141 | 556 | 420 | 39.7 |
| *D. rubicunda N* | DESeq2 | 498 | 253 | 245 | 300 | 39.7 |

**Table S3:** DEG recovery and gene annotation for EdgeR and DESeq2 analyses along with percent annotation using *Bombyx.* Up=FC>2, Implies night expression is greater than day expression. Down=FC<2, implies day expression is greater (control=Day, Night = Treatment)

| Species | Assembly annotation | BUSCO score | BUSCO duplication | No. of contigs* | Redundancy (/15k) |
|---|---|---|---|---|---|
| Dry_rub | Combined | 94.3 | 90.8 | 281790 | 18.8x |
| Dry_rub | Combined_cds_cdhit_95 | 90.1 | 80.2 | 184808 | 12.3x |
| Dry_rub | Combined_cds_filtered (v5) | 85.2 | 59.6 | 41712 | 2.8x |
| Dry_rub | Combined_cds_stringent_filter (v6) | 75.4 | 4.9 | 19877 | 1.3x |
| Ani_pel | Combined | 94.3 | 90.8 | 281790 | 18.78 |
| Ani_pel | Combined_cds | 93.9 | 90.3 | 233097 | 13.3x |
| Ani_pel | Combined_cds_filtered_cds (v5) | 85.2 | 59.2 | 41712 | 2.8x |
| Ani_pel | Combined_cds_stringent_filter (v6) | 79.3 | 4.7 | 19087 | 1.3x |

**Table S4:** BUSCO and contig distribution statistics for different species' *de-novo* assemblies with and without filtering. Increased filtering reduces redundancy and duplication but comes at the cost of reducing BUSCO scores. *All statistics are based on contigs of size >= 500 bp.

| nGenes | Fold Enrichment | Pathway | GO_ID | Gene Names |
|---|---|---|---|---|
| 2 | 85.53 | **Regulation of excretion** | GO:0044062 | NPHS1,SLC9A3R1 |

| | | | | |
|---|---|---|---|---|
| 1 | 42.76 | Response to herbicide | GO:0009635 | Alad |
| 1 | 42.76 | Carbohydrate catabolic process | GO:0016052 | BMSK0015853 |
| 1 | 42.76 | Blastocyst formation | GO:0001825 | Actl6a |
| 1 | 42.76 | Tryptophan catabolic process to kynurenine | GO:0019441 | KFase |
| 1 | 42.76 | Microvillus assembly | GO:0030033 | SLC9A3R1 |
| 1 | 42.76 | Parallel actin filament bundle assembly | GO:0030046 | jv |
| 1 | 42.76 | Heparan sulfate proteoglycan metabolic process | GO:0030201 | botv |
| 1 | 42.76 | Inactivation of MAPK activity | GO:0000188 | Spred1 |
| 1 | 42.76 | B cell activation | GO:0042113 | AKAP17A |
| 1 | 42.76 | Negative regulation of glycogen biosynthetic process | GO:0045719 | Gfpt1 |
| 1 | 42.76 | Behavioral response to pain | GO:0048266 | pain |
| 1 | 42.76 | Interphase microtubule nucleation by interphase microtubule organizing center | GO:0051415 | Grip91 |
| 1 | 28.51 | Cellular defense response | GO:0006968 | PNLIPRP2 |
| 1 | 28.51 | Cartilage condensation | GO:0001502 | COL11A1 |
| 1 | 28.51 | Short-chain fatty acid import | GO:0015913 | SLC5A8 |
| 1 | 28.51 | Male genitalia development | GO:0030539 | ken |
| 1 | 28.51 | Microtubule organizing center organization | GO:0031023 | Gcc2 |
| 1 | 28.51 | Telomere maintenance via semi-conservative replication | GO:0032201 | RFC3 |
| 1 | 28.51 | Glomerular basement membrane development | GO:0032836 | NPHS1 |
| 1 | 28.51 | Endosomal vesicle fusion | GO:0034058 | Ankfy1 |
| 1 | 28.51 | High-density lipoprotein particle assembly | GO:0034380 | ABCA7 |
| 1 | 28.51 | Positive regulation of Rho protein signal transduction | GO:0035025 | MCF2L |
| 1 | 28.51 | Xenobiotic transport | GO:0042908 | Mdr49 |
| 1 | 28.51 | DNA replication-removal of RNA primer | GO:0043137 | Rnaseh1 |
| 2 | 28.51 | Negative regulation of endocytosis | GO:0045806 | ABCA7,Mctp1 |
| 2 | 28.51 | **Negative gravitaxis** | GO:0048060 | pain,pyx |
| 1 | 28.51 | Imaginal disc-derived wing expansion | GO:0048526 | Spn88Ea |
| 1 | 28.51 | **Positive regulation of synapse structural plasticity** | GO:0051835 | FRMPD4 |
| 1 | 28.51 | Chitin biosynthetic process | GO:0006031 | chs-2 |
| 2 | 19.01 | Lysosomal transport | GO:0007041 | Vps53,g |
| 2 | 19.01 | RNA interference | GO:0016246 | Dcr-1,bel |

| | | | | |
|---|---|---|---|---|
| 2 | 15.55 | Negative regulation of endopeptidase activity | GO:0010951 | Spn88Ea,Serpinb1a |
| 2 | 13.16 | Formation of cytoplasmic translation initiation complex | GO:0001732 | eIF3-S5, BMSK0008517 |
| 2 | 10.69 | Protein heterotetramerization | GO:0051290 | pyx,Agrn |
| 2 | 10.06 | **Positive regulation of synaptic growth at neuromuscular junction** | GO:0045887 | Agrn,Sin1 |
| 2 | 8.55 | Endoplasmic reticulum organization | GO:0007029 | atl,Lman1 |
| 2 | 7.78 | **Regulation of synaptic growth at neuromuscular junction** | GO:0008582 | atl,hiw |
| 2 | 7.13 | Negative regulation of cell migration | GO:0030336 | SLC9A3R1,Mctp1 |
| 5 | 5.94 | **Visual perception** | GO:0007601 | RDH11,COL11A1, OP1,WDR36,disco |

**Table S5:** Top significantly enriched GO terms for *A. pellucida* day upregulation

| **No. Genes** | **Fold Enrichment** | **Pathway** | **GO_ID** | **Gene Names** |
|---|---|---|---|---|
| 1 | 41.61 | Female meiosis II | GO:0007147 | polo |
| 1 | 41.61 | Female germline ring canal formation-actin assembly | GO:0008302 | Src64B |
| 1 | 41.61 | Dosage compensation by hyperactivation of X chromosome | GO:0009047 | mle |
| 1 | 41.61 | 10-formyltetrahydrofolate catabolic process | GO:0009258 | aldh1l1 |
| 1 | 41.61 | **Sialic acid transport** | **GO:0015739** | **SLC17A5** |
| 1 | 41.61 | Arachidonic acid metabolic process | GO:0019369 | FAAH2 |
| 1 | 41.61 | Regulation of cellular metabolic process | GO:0031323 | melt |
| 1 | 41.61 | Protein exit from endoplasmic reticulum | GO:0032527 | SEC16A |
| 1 | 41.61 | Single strand break repair | GO:0000012 | gkt |
| 1 | 41.61 | Ring gland development | GO:0035271 | gl |
| 1 | 41.61 | Notch receptor processing-ligand-dependent | GO:0035333 | Nct |
| 1 | 41.61 | Positive regulation of RNA polymerase II transcriptional preinitiation complex assembly | GO:0045899 | PSMC6 |
| 1 | 41.61 | Tetrahydrofolate metabolic process | GO:0046653 | MTHFS |
| 1 | 41.61 | Neurotrophin TRK receptor signaling pathway | GO:0048011 | Bcar1 |
| 1 | 41.61 | Glucose catabolic process | GO:0006007 | agp |
| 1 | 41.61 | Glutaminyl-tRNAGln biosynthesis via transamidation | GO:0070681 | gatA |
| 1 | 41.61 | Error-free translesion synthesis | GO:0070987 | POLH |
| 1 | 41.61 | Protein deubiquitination involved in ubiquitin-dependent protein catabolic process | GO:0071947 | trbd |

| | | | | |
|---|---|---|---|---|
| 1 | 41.61 | Negative regulation of release of cytochrome c from mitochondria | GO:0090201 | Parl |
| 1 | 41.61 | Regulation of cardiac conduction | GO:1903779 | nkx-2.5 |
| 1 | 41.61 | Methionyl-tRNA aminoacylation | GO:0006431 | MetRS-m |
| 2 | 33.29 | Tetrahydrofolate interconversion | GO:0035999 | Sardh,MTHFS |
| 1 | 27.74 | Tubulin complex assembly | GO:0007021 | Tbcd |
| 1 | 27.74 | Double-strand break repair via break-induced replication | GO:0000727 | Cdc45 |
| 1 | 27.74 | Aminophospholipid transport | GO:0015917 | ATP11B |
| 1 | 27.74 | **Synaptic target attraction** | **GO:0016200** | **Ten-a** |
| 1 | 27.74 | N-terminal peptidyl-methionine acetylation | GO:0017196 | Naa60 |
| 1 | 27.74 | Peptidyl-lysine methylation | GO:0018022 | Eef1akmt2 |
| 1 | 27.74 | Negative regulation of ossification | GO:0030279 | LRP4 |
| 1 | 27.74 | Negative regulation of phosphoprotein phosphatase activity | GO:0032515 | Bod1 |
| 1 | 27.74 | Store-operated calcium entry | GO:0002115 | olf186-F |
| 1 | 27.74 | Cytoplasmic sequestering of transcription factor | GO:0042994 | Sufu |
| 1 | 27.74 | Melanin biosynthetic process from tyrosine | GO:0006583 | BMSK0009475 |
| 1 | 20.80 | Cytoplasmic mRNA processing body assembly | GO:0033962 | patl1 |
| 1 | 20.80 | Positive regulation of transcription from RNA polymerase I promoter | GO:0045943 | HEATR1 |
| 1 | 20.80 | Negative regulation of insulin secretion | GO:0046676 | HADH |
| 1 | 16.64 | Microtubule nucleation | GO:0007020 | TUBG1 |
| 1 | 16.64 | **Eclosion rhythm** | **GO:0008062** | **disco** |
| 1 | 16.64 | Late endosome to vacuole transport | GO:0045324 | chmp3 |
| 1 | 16.64 | **Negative regulation of axon extension involved in axon guidance** | **GO:0048843** | **tap** |
| 1 | 16.64 | Glucose transmembrane transport | GO:1904659 | SLC2A3 |
| 1 | 16.64 | Nucleocytoplasmic transport | GO:0006913 | Anp32a |
| 2 | 13.87 | Chromosome segregation | GO:0007059 | PPP2R1A,Naa60 |
| 1 | 13.87 | Nucleotide catabolic process | GO:0009166 | BMSK0009399 |
| 1 | 13.87 | Phosphatidylethanolamine biosynthetic process | GO:0006646 | PCYT2 |
| 2 | 12.80 | Negative regulation of sequence-specific DNA binding transcription factor activity | GO:0043433 | melt,Sufu |
| 2 | 10.40 | Protein heterotetramerization | GO:0051290 | pyx,LRP4 |
| 2 | 9.25 | Phospholipid biosynthetic process | GO:0008654 | Mboat1,PCYT2 |

| | | | | |
|---|---|---|---|---|
| 2 | 9.25 | **Locomotor rhythm** | **GO:0045475** | **Ork1,disco** |
| 2 | 8.76 | Pupal chitin-based cuticle development | GO:0008364 | Gld,Gld |
| 3 | 8.61 | Mitochondrial translation | GO:0032543 | BMSK0007759,MRPS35,gatA |
| 2 | 8.32 | Sperm storage | GO:0046693 | Gld,Gld |
| 2 | 7.93 | Cytoplasmic translation | GO:0002181 | RpL4, |
| 2 | 7.93 | Synaptic growth at neuromuscular junction | GO:0051124 | LRP4,Ten-a |
| 2 | 7.56 | Negative regulation of canonical Wnt signaling pathway | GO:0090090 | PSMC6,LRP4 |
| 2 | 6.93 | **Mushroom body development** | **GO:0016319** | **tap,Src64B** |
| 2 | 6.93 | Post-translational protein modification | GO:0043687 | TULP4,PSMC6 |
| 2 | 6.93 | **Synapse organization** | **GO:0050808** | **Ten-a,LRP4** |
| 2 | 5.74 | **Trehalose transport** | **GO:0015771** | **Tret1-1,Tret1-1** |
| 2 | 5.20 | Positive regulation of canonical Wnt signaling pathway | GO:0090263 | PSMC6,trbd |
| 3 | 5.09 | Cell migration | GO:0016477 | trbd,Bcar1,SPEF1 |
| 2 | 5.04 | Response to endoplasmic reticulum stress | GO:0034976 | SEC16A,Pdi |
| 3 | 4.62 | Mitotic cell cycle | GO:0000278 | TUBG1,Tbcd,polo |
| 2 | 4.62 | Microtubule cytoskeleton organization | GO:0000226 | TUBG1,Tbcd |
| 3 | 3.37 | G-protein coupled receptor signaling pathway | GO:0007186 | Bcar1,Atrnl1,TRHR |

**Table S6:** Top significantly enriched GO terms for *D. rubicunda* night upregulation

| Gene Symbol | Gene Name | Function | DESeq2 | EdgeR | WGCNA | Flybase ID | SILKD ID |
|---|---|---|---|---|---|---|---|
| BNC1/disco | disconnected | vision | 1* | 1* | | FBgn0000459 | BMSK0012241 |
| Spt5 | Spt5 | transcription | 1* | 1* | | FBgn0040273 | BMSK0011143 |
| 1q/trbd | trabid | development | | 1 | | FBgn0037734 | BMSK0015294 |
| RpL4 | Ribosomal protein L4 | energy use, Ribosomal production | 1* | | | FBgn0003279 | BMSK0006059 |
| Prp2/PPO2 | Prophenoloxidase 2 | Wound melanization | 1* | | | FBgn0033367 | BMSK0009475 |
| mRpS5 | mitochondrial ribosomal proteins | energy use, mitochondrial maintenance | 1* | | | FBgn0287187 | BMSK0014662 |
| SLC2A6/Tret1-1 | Solute Carrier Family 2 Member 6 | locomotion, energy metabolism | 1* | | | FBgn0050035 | BMSK0003817**, BMSK0003818 |

| | | | | | | | |
|---|---|---|---|---|---|---|---|
| TUBG1/γTub 23C | tubulin gamma 1 | brain development in adult | 1* | | | FBgn0260639 | BMSK0002451 |
| PARL/rho-7 | presenilin associated rhomboid like | mitochondrial maintenance | 1* | | | FBgn0033672 | BMSK0008635 |
| SLC17A5/MFS10 | Solute carrier family 17 member 5 | transmembrane ion transporter | 1* | | | FBgn0030452 | BMSK0001219 |
| titin1/sls | sallimus | locomotion | 1 | | | FBgn0086906 | BMSK0000202 |
| unc-22/unc | uncoordinated | adult locomotion, hearing | 1 | | 1 | FBgn0003950 | BMSK0015609**, BMSK0000066 |
| ARHGAP17/RhoGAP92B | rho GTPase-activating protein 17 | auditory, brain development | | | 1 | FBgn0038747 | BMSK0009977 |
| tk | tachykinin | locomotion and circadian rhythms | | | 1 | FBgn0037976 | BMSK0003725 |
| BmorOBP1/Obp58a | Odorant Binding Protein 1 | odor binding | | | 1 | FBgn0034768 | BMSK0013390 |
| PPAP2A/wun | Inorganic Pyrophosphatase 2/ wunnen | vision | | | 1 | FBgn0016078 | BMSK0011305 |
| BmorOBP2/Obp84a | Odorant Binding Protein 2 | odor binding | | | 1 | FBgn0011282 | BMSK0009610 |
| takt/JHBP | takeout-like | circadian | | | 1 | FBgn0038395 | BMSK0013046 |

**Table S7:** Gene symbol from EggNog or *B. mori* annotation. Gene names, function and closest flybase ID orthologs used to collect evidence for the various listed functions.* Indicated that the gene altered direction or trend of expression across a diel pair. ** Indicates this ortholog was used for structural modeling and conservation analyses. This list is only a subset of the genes recovered from various analyses. The genes were chosen because they had either 1) multiple lines of evidence, robust divergent expression pattern, or 2) a gene ontology term for sensory or circadian function among three datasets. For exhaustive lists, see Supplementary Data

| Process | Search Query | Excluding |
|---|---|---|
| Vision | "vision" or "eye" or "visual" or " compound eye" or "visual perception" or "light" or "ocellus" or "visual" or "photoreceptor" | - "chain" - "immunoglobulin" - "light harvesting" - "photosynthesis" - "plastid" -"plant" -actin -myosin -photoperiod - touch -smell -"light chain" |

| | | |
|---|---|---|
| Smell | "olfaction" or "smell" or "antennal" or "olfactory" or "odor" or "pheromone" | |
| Hearing | "audition" or "hearing" or "sound" or "auditory" or "tympanum" or "ear" | |
| Circadian | "circadian" or "rhythm" or "clock" or "diurnal" or "nocturnal" or "entrainment" | |
| Behaviour | "locomotion" or "flight" or "behavioral" | - "cell" - "morphogenesis" - "development" or - "pheromone" |
| Brain | "neural plasticity" or "brain" or "neuropeptide" | |

**Table S8:** GO query terms for functional lookup

### Legends for Datasets

Supp_dataset_1: EdgeR: EdgeR sample metadata, analysis parameters (config files for RasFlow), differentially expressed gene sets, annotated DEG sets with *Bombyx mori* annotations for *Anisota pellucida* (Ap) and *Dryocampa rubicunda* (Dr). Overlapping genes between *Anisota* and *Dryocampa.* Analysis performed with de-novo assembly versions 5 for both species.

Supp_dataset_2: DESeq2: DESeq2 sample metadata, analysis parameters (R script), differentially expressed gene sets, and annotated DEG sets with *Bombyx mori* annotations for *Anisota pellucida* (Ap) and *Dryocampa rubicunda* (Dr). Unique and overlapping genes between *Anisota* and *Dryocampa.* Analysis performed with de-novo assembly versions 5 for both species.

Supp_dataset_3: Supporting_Table_Common_RNAseq: Overlapping genes for both analyses with a pivot table summary of the different unique *Bombyx* genes and the number of transcripts mapped to each.

Supp_dataset_4: Gene_modules_WGCNA: WGCNA identified modules for *Anisota* and *Dryocampa* individual count data and combined modules along with annotations of the (grey60, tan, turquoise and blue) modules.

Supp_dataset_5: GO_analyses: TopGO, ShinyGo and Revigo analyses

Supp_dataset_6:Analyses_combined:All analyses results combined and annotated, with FC<0 and FC<2 for DEGs. Note to combine analyses fold change signs were switched for EdgeR changed.

Supp_dataset_7: Overlap_common_genes_annotated_with_sequence: Overlapping transcripts annotated with sequences for both species and *Bombyx mori.* Note to combine analyses fold change signs were switched for EdgeR changed.

Supp_dataset_8_EggNOG_annotations: EggNog annotations of the various sequences in dataset 7 and the GO terms used to query this dataset

Supp_datatset_9: GO_lookup_genes: Genes recovered from the GO cross referencing with EggNOG annotations

Supp_dataset_10: Genes of interest for which conservation and protein models were constructed

Supp_dataset_11: Assembly_codes: 11a:List of moths and assembly codes used 11b: List of insects and assembly codes used for Orthofinder searches

Supp_dataset_12_Alphafold_Bmor_models: Alphafold predicted models for *Bombyx* genes of interest

Supp_dataset_13_Conservation analyses: Consurf predicted models and conservation analyses for insects and moths

Supp_dataset_14_PyMOL: PyMOL files showcasing the overlapping protein structures
